# TOX enforces the immunosuppressive program of tumor-infiltrating regulatory T cells

**DOI:** 10.64898/2026.06.19.732838

**Authors:** Seyeon Park, Dong Jin Park, Myeong Joon Kim, Geoffrey Kelly, Ruihua Zhang, Gamin Kim, Joon Jeong, Seunghee Kim-Schulze, Hye Ryun Kim, Kyungsoo Kim, Sang-Jun Ha

**Author notes:** Correspondence to: Sang-Jun Ha, Department of Biochemistry, College of Life Science & Biotechnology, Yonsei University, Seoul 03722, Korea., Kyungsoo Kim, Institute for Breast Cancer Precision Medicine, Yonsei University College of Medicine, Seoul 06229, Korea.

## Abstract

Regulatory T (Treg) cells accumulate in the tumor microenvironment (TME) to suppress anti-tumor immunity, but the transcriptional regulators stabilizing their immunosuppressive state remain poorly defined. Here we show that transcriptional factor TOX is selectively upregulated in tumor-infiltrating (TI) Treg cells across human cancers and mouse tumor models, while remaining low in peripheral and naive Treg populations. Treg-specific deletion of TOX reduced tumor burden, impaired TI Treg-mediated immune suppression, and enhanced effector functions of CD8⁺ and CD4⁺ T cells. In mosaic mice, TOX-deficient Tregs were selectively depleted from tumors, accompanied by increased apoptosis. Single-cell RNA sequencing and TCR clonotype analysis linked TOX expression to an effector-like TI Treg state with clonal expansion, whereas TOX loss shifted cells toward a TCF7-associated progenitor-like phenotype. ATAC-seq revealed enrichment of AP-1 motifs in TOX-sufficient TI Tregs. In contrast, TCF7, LEF1, and FOXO1 motifs in TOX-deficient counterparts, uncovering the opposing transcriptional networks downstream of TOX. Furthermore, TOX deficiency augmented CD8⁺ T cell responses to PD-1 blockade. Together, these findings establish TOX as a key regulator of TI Treg fitness and stability, and identify it as a potential therapeutic target to enhance the efficacy of PD-1-based immunotherapy.

## INTRODUCTION

Tumor progression is accompanied by sustained immune activation in the tumor microenvironment (TME), yet antitumor immunity frequently collapses under dominant immunosuppressive programs. Regulatory T (Treg) cells, defined by transcription factor Foxp3, are indispensable for immune homeostasis and prevention of autoimmunity,^1, 2^ but in cancer they accumulate in the TME and enforce potent suppression that limits protective T cell immunity and supports tumor growth.^3, 4^ Consequently, elevated intratumoral Treg abundance, or an increased Treg-to-CD8^+^ T cell ratio, correlates with poor clinical outcome and reduced response to immune checkpoint blockade across multiple tumor types.^4, 5, 6^ Although targeting Tregs shows therapeutic promise, systemic Treg depletion carries substantial risk of autoimmunity and immune-related Adverse Events (irAEs). This underscores the need to identify tumor-specific Treg programs that can be selectively disrupted without compromising peripheral immune tolerance.^7, 8^

A critical feature distinguishing tumor-infiltrating (TI) Tregs from their peripheral counterparts is the acquisition of an effector-like state characterized by high expression of checkpoint receptors - including CTLA-4, PD-1, TIGIT, and ICOS - as well as enhanced production of suppressive molecules such as IL-10, TGF-β, and granzyme B.^9, 10, 11, 12^ This effector TI Treg phenotype is associated with superior suppressive capacity compare to peripheral Tregs and is maintained even under the inflammatory conditions of the TME, suggesting active transcriptional reinforcement rather than passive accumulation.^13, 14^ While transcription factors such as BATF, BLIMP-1, IRF4, and HIF-1α have been implicated in aspects of Treg effector differentiation or adaptation to metabolic stress,^15, 16, 17, 18, 19^ none has emerged as a tumor-context–specific inducible regulators. Specifically, a factor that comprehensively stabilizes the checkpoint-high, effector TI Treg phenotype in the TME while remaining dispensable for systemic Treg homeostasis has yet to be identified. Uncovering such a regulator represents a central unmet need in the field.

TOX is a high-mobility-group (HMG) box transcription factor with diverse roles in immune development and differentiation,^20, 21, 22, 23, 24, 25, 26, 27^ and it is best known as a key driver of CD8^+^ T cell exhaustion in chronic infection and cancer.^28, 29, 30, 31, 32, 33, 34^ Persistent antigen stimulation induces TOX expression, enabling widespread chromatin remodeling and transcriptional stabilization of exhaustion-associated programs in CD8^+^ T cells.^29, 31^ High TOX expression in TI CD8^+^ T cells correlates with poor prognosis,^30, 31, 32, 33, 35, 36^ and our prior work identified TOX as a driver of the exhaustion program in human TI CD8^+^ T cells.^30^ This established role raises a fundamental conceptual question: whether TOX, which enforces a terminally dysfunctional, exhausted state in CD8^+^ T cells, might paradoxically stabilize an active suppressive effector program in a distinct intratumoral lymphocyte lineage, namely Tregs. This possibility is supported by emerging observations: TOX has been detected in Tregs in human malignancies including non-Hodgkin lymphoma^37^ and multiple myeloma,^34^ and is induced in tonic-signaling CAR-engineered human Treg cells in association with functional remodeling.^38^ However, whether TOX actively regulates TI Treg identity and suppressive capacity in the TME - rather than merely reflecting a generalized activation or chronic stimulation signal - has remained unknown. Equally unresolved is whether TOX operates as a constitutive lineage-associated factor in Tregs broadly, or as an inducible regulator selectively engaged by cues unique to the TME.

Here, we show that TOX is an inducible transcriptional and epigenetic regulator that is selectively engaged in TI Tregs across human cancers and mouse tumor models, and that it is required for the acquisition and maintenance of an effector-like, checkpoint-high suppressive program in the TME. Treg-specific deletion of TOX reduces tumor burden and unleashes robust CD4⁺ and CD8⁺ T cell effector responses in the TME. Within-host mosaic analyses establish that TOX-deficient TI Tregs are cell-intrinsically outcompeted, lose effector-associated features, and undergo apoptosis in the same tumor environment that supports wild-type TI Tregs. Integrated single-cell transcriptomic, V(D)J clonotype, and chromatin accessibility profiling reveals that TOX stabilizes an AP-1–driven, clonally expanded effector TI Treg state, while its loss redirects TI Tregs toward a less differentiated TCF7-associated program. Furthermore, destabilization of the TOX-dependent TI Treg program markedly enhances the efficacy of PD-1 blockade. Together, these findings establish TOX as a tumor-context–specific regulator that enforces immunosuppressive identity of intratumoral Treg, providing a compelling mechanistic rationale for targeting TOX-dependent circuitry to selectively disrupt intratumoral suppression and potentiate antitumor immunity.

## METHODS

### Human samples for mass cytometric analysis (CyTOF)

Human tumor samples were obtained from patients with head and neck squamous cell carcinoma (HNSCC, *n* = 23) and non-small cell lung carcinoma (NSCLC, *n* = 35) undergoing surgical resection at Severance Hospital, following informed consent (**Supplementary Table 1, 2**). The study was approved by the Institutional Review Board of Severance Hospital (IRB no. 4-2020-0945 and no. 4-2022-0130), and conducted in accordance with the Declaration of Helsinki.

Tumor tissues were mechanically dissociated into ∼1 mm³ fragments and enzymatically digested using the Tumor Dissociation Kit (Miltenyi Biotec), following the manufacturer’s instructions. Single-cell suspensions were stained with a panel of metal isotope–conjugated antibodies for CyTOF analysis. Antibodies were purchased either pre-conjugated or conjugated in-house with heavy metals using the MaxPar X8 Antibody Labeling Kit (Standard BioTools). To reduce nonspecific binding, cells were incubated with Human TruStain FcX^TM^ (BioLegend) prior to staining with surface antibodies. The viability dye 103Rh and Cell-ID^TM^ Intercalator-Ir reagents (191Ir, 193Ir) were included. Surface markers included CD45 (89Y, HI30), ICOS (151Eu, C398.4A), and OX40 (158Gd, ACT35) (all from Standard BioTools); TIM-3 (156Gd, A18087E), CD25 (166Er, BC96), and PD-1 (175Lu, EH12.2H7) (all from BioLegend); CD19 (142Nd, REA675), CD4 (145Nd, REA623), CD8 (146Nd, REA734), CD66b (152Sm, REA306), TIGIT (153Eu, REA1004), CD56 (161Dy, REA196), and CD3 (168Er, REA613) (all from Miltenyi Biotec); and CD127 (149Sm, A019D5) (from Fluidigm). Cells were subsequently fixed and permeabilized using the Foxp3 Fixation/Permeabilization Kit (eBioscience), followed by intracellular staining with CTLA-4 (162Dy, 14D3) (from Standard BioTools), CD68 (115In, Y1/82A) (from BioLegend), TOX (171Yb, REA473) (from Miltenyi Biotec), and Foxp3 (159Tb, PCH101) (from Fluidigm). Data acquisition was performed on a Helios mass cytometer (Standard BioTools). Signal normalization across samples was achieved using the bead-based normalization method. For quality control, CD45^+^ populations were gated in FlowJo software to exclude debris, dead cells, and doublets, retaining only live, single immune cells for analysis. Following stringent data cleaning, CyTOF data were processed using R. T cells (CD3^+^) were identified after exclusion of B cells (CD3^−^ CD19^+^), granulocytes (CD66b^+^), macrophages (CD3^−^ CD68^+^), and NK cells (CD3^−^ CD56^+^). Within the T cell compartment, subsets were defined as CD8^+^ T cells (CD3^+^ CD8^+^), conventional CD4^+^ T cells (CD3^+^ CD4^+^ Foxp3^−^), and regulatory T cells (CD3^+^ CD4^+^ Foxp3^+^ CD25^+^ CD127^−^).

### Mice

C57BL/6N mice were purchased from the Jackson Laboratory. The C57BL/6N *Tox*^tm1a(KOMP)Wtsi^ mice (KOMP ES cell line; RRID:MMRRC_064029-UCD) were obtained from the Mutant Mouse Resource and Research Center (MMRRC) at the University of California, Davis, an NIH-funded strain repository (PMID: 21677750). The *Tox*^tm1a(KOMP)Wtsi^ mice were crossed with *Actin*-Flippase mice to excise the FRT-flanked neomycin resistance cassette, generating *Tox*^tm1c(KOMP)Wtsi^ mice (hereafter referred to as *Tox*^fl/fl^). *Actin*-Flippase mice were generously provided by Hyoung Pyo Kim (Yonsei University). *Tox*^fl/fl^ mice were further crossed with *Foxp3*^YFP-Cre^ mice, which were kindly provided by Yeonseok Chung (Seoul National University), to generate TOX conditional knockout mice. All mice were maintained under specific pathogen-free conditions at Yonsei University. Six- to eight-week-old male or female mice were used for all experiments. All animal procedures were reviewed and approved by the Institutional Animal Care and Use Committee (IACUC) of Yonsei University (IACUC-A-202111-1369-01 and IACUC-A-202412-1979-01).

### *In vivo* tumor models

The murine CT26 and MC38 colon adenocarcinoma, TC-1 lung adenocarcinoma, and B16F10 melanoma cell lines were purchased from ATCC in 2010. Cells were subjected to annual Mycoplasma testing using the e-Myco Mycoplasma PCR Detection Kit (iNtRON Biotechnology). All murine tumor cells were cultured in RPMI or DMEM supplemented with 10% FBS, penicillin (100 U/ml), and streptomycin (100 mg/ml) for two passages prior to *in vivo* injection. A total of 5 × 10^5^ tumor cells were injected either intravenously via the tail vein or subcutaneously into C57BL/6N mice as well as various genetically modified mouse strains including Tox^fl/fl^ (TOX^Ctrl^), Tox^fl/fl^ × Foxp3^YFP-Cre (+/+)^ (TOX^cKO^), and Tox^fl/fl^ × Foxp3^YFP-Cre (+/-)^ mice (TOX^cre+/-^). Mice were euthanized on day 20 post-tumor injection, and spleen, lung, or tumor tissues were harvested for single-cell suspension preparation. For the staggered injection, wild-type mice were inoculated with TC-1 tumor cells on days 7, 14, and 20, with all mice subsequently euthanized simultaneously. For the survival study, TC-1 tumor-bearing TOX^Ctrl^ or TOX^cKO^ mice were observed until mortality or until severe distress necessitated euthanasia.

### LCMV chronic infection model

C57BL/6N, TOX^Ctrl^, and TOX^cKO^ mice were intravenously infected with LCMV Clone 13 (Cl13) at a concentration of 2 × 10^6^ plaque-forming units (p.f.u.) in serum-free RPMI medium. LCMV-Cl13, a derivative originating from an LCMV Armstrong (Arm) AC1371 host mouse, was acquired from Rafi Ahmed (Emory Vaccine Center). To assess degranulation and intracellular cytokine production in T cells, splenocytes were stimulated with LCMV-glycoprotein peptides (GP), including 0.2 μg/ml of GP33–41 and GP276–286, and a peptide pool containing 5.0 μg/ml of GP66–80, 0.2 μg/ml of GP70–77, GP92–101, and GP118–125, in the presence of GolgiPlug/GolgiStop at 37°C for 5 hours. Cells were subsequently stained for intracellular cytokines, including IFN-γ, TNF, IL-2, and CD107a. Intracellular staining was conducted using the BD Cytofix/Cytoperm fixation and permeabilization kit (BD Biosciences) according to the manufacturer’s protocol.

### Isolation of lymphocytes

Lymphocytes were isolated from the spleen, lung, or tumor tissue in wild-type, tumor-bearing mice or virus-infected mice. The spleens were minced onto a 40-μm cell strainer (Falcon, Corning), and red blood cells were removed using ACK lysis buffer (Gibco). The lung or tumor tissues were minced into 1mm^3^ fragments and then digested with 1 mg/mL collagenase type IV (Worthington Biochemical Corp.) and 0.01 mg/mL DNase I (Millipore Sigma Corp.) in 10% FBS containing RPMI media at 37 °C for 30 mins. The dissociated tissues were filtered using a 40-μm cell strainer (Falcon, Corning), and red blood cells were removed using ACK lysis buffer. The single-cell suspensions were stained with the indicated fluorescent dye-conjugated antibodies.

### Antibodies and flow cytometry

The single-cell suspensions were stained with the following fluorochrome-conjugated antibodies in PBS supplemented with 2% FBS: CD4 (RM4-5), CD8 (53-6.7), CD25 (PC61), CD39 (Duha59), CD44 (IM7), CD69 (H1.2F3), CD45.2 (104), CD127 (A7R34), PD-1 (29F.1A12), TIM-3 (RMT3-23), LAG (C9B7W), TIGIT (1G9), ICOS (C398.4A), KLRG1 (2F1/KLRG1), TNF-α (MP6-XT22), CCR4 (2G12), CCR8 (SA214G2) (all from BioLegend); CD8 (53-6.7), CD25 (PC61.5), CD103 (2E7), CTLA-4 (UC10-4B9), Foxp3 (FJK-16s) (all from Invitrogen); CD107a (1D4B), Ki67 (B56), IFN-γ (XMG1.2), IL-2 (JES6-5H4) (all from BD biosciences); and TOX (REA473) (from Miltenyi Biotec). Dead cells were excluded by staining the LIVE/DEAD Fixable Near-IR Dead Cell Stain Kit (Invitrogen). For intracellular staining, cells were fixed and permeabilized using the Foxp3 fixation/permeabilization solution (eBioscience, Cat#00-5521-00), followed by staining for Foxp3, CTLA-4 and Ki67. For cytokine staining, cells were fixed and permeabilized using BD Cytofix/Cytoperm fixation/permeabilization solution (BD Biosciences, Cat#554722), followed by staining for IFN-γ, TNF-α and IL-2. To detect the expression of intracellular cytokines, lymphocytes from the spleen and tumor tissue were stimulated with phorbol 12-myristate 13-acetate (PMA, 50 ng/ml) and ionomycin (500 ng/ml) in the presence of GolgiStop (BD Biosciences, Cat#554724) and GolgiPlug (BD Biosciences, Cat#555029) at 37°C for 6 hours. Flow cytometry analysis was performed using the CytoFLEX or CytoFLEX LX (Beckman Coulter). All data were analyzed using FlowJo software (Tree Star).

### *Ex vivo* suppression assay

For the *ex vivo* Treg suppression assay, Tregs were isolated from TC-1 tumor-bearing TOX^Ctrl^ or TOX^cKO^ mice using CD4^+^CD25^+^ Regulatory T Cell Isolation kit (Miltenyi Biotec), respectively. CD8^+^ T cells were isolated from the naïve mouse using CD8^+^ T Cell Isolation kit (Miltenyi Biotec). Following the isolation of splenic CD8^+^ T cells, the cells were labeled with CellTrace™ Violet (CTV, Thermo Fisher Scientific) in accordance with the manufacturer’s instructions. Tregs derived from TOX^Ctrl^ or TOX^cKO^ mice were co-cultured with CTV-labeled CD8^+^ T cells in the presence of Dynabeads™ mouse T-activator CD3/CD28 (Thermo Fisher Scientific) for 72 hours at different ratio. The proliferation of CD8^+^ T cells was measured by flow cytometry (CytoFLEX LX, Beckman Coulter). To detect the secretion of IFN-γ, CTV-labeled CD8^+^ T cells were re-stimulated with soluble anti-CD3 (2μg/ml) and anti-CD28 (2μg/ml) antibodies, supplemented with GolgiPlug/GolgiStop for the final 6 hours of culture.

### Cell sorting

For flow cytometry analysis and single-cell RNA and V(D)J sequencing, spleens or tumors were harvested from TC-1 tumor-bearing TOX^cre+/−^ mice. The spleen and tumor were dissociated as previously described. Subsequently, the spleen or tumor samples were pooled and isolated CD4^+^ T cells using CD4^+^ T cell Isolation kit according to the manufacturer’s protocol (Miltenyi Biotec). The isolated CD4^+^ T cells were then stained with anti-CD4, anti-CD25, anti-CD127 antibodies, and the LIVE/DEAD Fixable Near-IR Dead Cell Stain kit. The stained cells were sorted into live YFP^−^CD25^+^ and YFP^+^CD127^−^CD4^+^ T cells for the flow cytometry analysis or into live YFP^−^CD25^−^, YFP^−^CD25^+^ and YFP^+^CD127^−^CD4^+^ T cells for single-cell RNA and V(D)J sequencing using BD FACS Aria^TM^ Fusion (BD Biosciences). For bulk ATAC sequencing, pooled tumor samples from TC-1 tumor-bearing TOX^Ctrl^ and TOX^cKO^ mice were dissociated as previously described. Tregs were sorted from each group into CD25^+^CD127^−^ or YFP^+^CD127^−^CD4^+^ T cell populations using the BD FACS Aria™ III (BD Biosciences).

### Apoptosis assay

To measure the apoptotic rate, lymphocytes from the spleen and tumor were isolated from TC-1 tumor-bearing TOX^cre+/−^ mice, as previously described. Subsequently, lymphocytes were stained using the BD Pharmingen™ PE Annexin V Apoptosis Detection Kit I, following the manufacturer’s protocol (BD Biosciences).

### Single-cell RNA and V(D)J sequencing analysis

Tumor samples were prepared as described above for single-cell RNA and V(D)J sequencing. The sorted live YFP^+^CD127^−^CD4^+^ T cells were stained with TotalSeq™-C0301 anti-mouse Hashtag 1 Antibody (Barcode Sequence: ACCCACCAGTAAGAC, BioLegend). Concurrently, the live YFP^−^CD25^−^ and YFP^−^CD25^+^CD127^−^CD4^+^ T cells were individually stained with TotalSeq™-C0302 anti-mouse Hashtag 2 Antibody (Barcode Sequence: GGTCGAGAGCATTCA, BioLegend) to optimize and enhance overall performance in accordance with the manufacturer’s instructions.

Single-cell sequencing library was prepared using the Chromium 10X Genomics platform with a Next GEM Single Cell 5. v2 (CG000330) according to the manufacturer’s instructions. Following dilution in nuclease-free water, single-cell suspensions were combined with a master mix and loaded onto a Next GEM Chip K along with Single Cell 5′ Gel Beads and Partitioning Oil. RNA transcripts were uniquely barcoded and reverse-transcribed in droplets. The resulting cDNA molecules were pooled and enriched via PCR, followed by size-based separation for 5ʹ Gene Expression libraries, V(D)J Enriched Libraries, and Cell Surface Protein libraries. For Cell Surface Protein Library, the cDNA pool was enriched with index PCR. For V(D)J Enriched Library, the cDNA enrichment was amplified twice with the TCR-specific primers. For 5’ Gene Expression Library, the cDNA pool underwent end repair, addition of a single ’A’ base, adapter ligation, purification, and PCR enrichment. Quantification was conducted using the KAPA qPCR Quantification Protocol, and qualification was carried out using the Agilent Technologies 4200 TapeStation. The libraries for both single-cell RNA sequencing (scRNA-seq) and single-cell V(D)J sequencing (scV(D)J-seq) were performed on the HiSeq platform (Illumina).

To process the scRNA-seq and scV(D)J-seq data, we used the Seurat R package and identified cells using designated hashtag antibodies C0301 and C0302. We demonstrated dimension reductions and identified Tregs using canonical markers including Foxp3. We reanalyzed dimension reductions using those Tregs. Gene expression in UMAP spaces were visualized using the FeaturePlot function in Seurat. Differentially expressed genes were identified using the FindMarkers function in the Seurat package. For V(D)J-seq data, we defined clonotypes assigned to multiple cells as “expanded clonotypes” and those assigned to a single cell as “unique clonotypes”. Gene set variation analysis (GSVA) was performed using the GSVA function in R, using log-transformed expression values from the Seurat object to calculate GSVA scores. Gene sets for GSVA were obtained from our previous study and MSigDB. Publicly available scRNA-seq data (GSE152022) were downloaded from the Gene Expression Omnibus (GEO) and analyzed using the Seurat package. Two datasets (GSM4598898, GSM4598899) were integrated using 2000 anchor features. Data visualization was performed using the DotPlot and VlnPlot functions within Seurat. For integration analysis, previously published our single-cell RNA sequencing data (GSE164033) were utilized. Tregs treated with isotype control were extracted from this dataset and integrated with our dataset except Mki67^+^ cluster 3 using the Seurat package.

GSEA was performed by using the function GSEA from the package clusterProfiler.^39^ The ranked gene list was imported from the differentially expressed genes calculated by comparing TOXKO versus TOXWT Tregs. The gene sets for GSEA were obtained from MSigDB^40, 41^ and a previous study.^42^

Differential expression between WT_1 and each knockout cluster (PD1_KO or TOX_KO) was computed using Seurat FindMarkers^43^ with logfc.threshold = 0 and min.pct = 0, and significance was defined as adjusted P < 0.05 (with an additional |avg_log2FC| > 0.25 flag recorded). Genes upregulated in WT_1 relative to PD1_KO and TOX_KO (avg_log2FC > 0, adjusted P < 0.05) were collected as WTHigh gene sets. Overlap between the two WTHigh sets was visualized as a Venn diagram using the ggvenn R package.

GSVA^44^ enrichment scores were computed (GSVA R package) on Seurat integrated expression data for cells restricted to WT_1 vs PD1_KO and WT_1 vs TOX_KO, using MSigDB gene sets (GS.MSigDB).^40, 41^ For each gene set, WT–KO differences were defined as the mean GSVA score in WT_1 minus the mean score in the corresponding KO cluster, and significance was assessed by two-sided Wilcoxon rank-sum tests followed by multiple-testing correction (Benjamini–Hochberg; adjusted P < 0.05). Gene sets were categorized as PD1_WTHigh, TOX_WTHigh, or Double_WTHigh when the WT–KO difference was positive and significant in the respective comparison(s), and PD1_Diff vs TOX_Diff values were visualized in a scatter “volcano-like” plot with significant terms labeled.

Scaled expression values from the Seurat integrated assay were extracted for WT-high DEGs, defined as genes upregulated in WT_1 versus PD1_KO and/or TOX_KO cells. Genes were grouped into PD1KO-specific, shared, or TOXKO-specific categories, and cells were restricted to WT_1, TOX_KO, and PD1_KO clusters. Mean scaled expression was calculated per cluster for each gene and visualized using pheatmap without hierarchical clustering, applying a green–black–red color scale with expression values capped at −2 to 2.

### Bulk ATAC sequencing analysis

To isolate nuclei, 1 × 10^5^ cells were prepared and lysed in cold lysis buffer. Nuclei concentration was quantified using the LUNA-FL™ Automated Fluorescence Cell Counter (Logos Biosystems), and morphology was assessed by microscopy. Immediately post-lysis, nuclei (5 × 10^4^ cells) were resuspended in a transposition reaction mix, prepared with the Tagment DNA TDE1 Enzyme and Buffer Kit (Illumina). The transposition reaction was incubated at 37°C for 30 minutes. Following transposition, DNA was purified using the MinElute PCR Purification Kit (Qiagen). Transposed DNA fragments were amplified with the optimal PCR cycle number determined by quantitative PCR (qPCR) to minimize GC and size bias. A side reaction qPCR was run to calculate the additional cycles required, plotting linear Rn versus cycle to identify the cycle number at 1/4 maximum fluorescence intensity. The remaining PCR reaction was run to this cycle threshold. Purified libraries were subsequently quantified by qPCR following the KAPA qPCR Quantification Protocol and qualified with a Bioanalyzer (Agilent Technologies). Finally, libraries were sequenced on an Illumina HiSeq or NovaSeq platform.

Genome-wide ATAC-seq peaks (MACS2^45^ broad peaks filtered to remove ENCODE blacklist regions^46^) were annotated using ChIPseeker^47^ annotatePeak with a ±3 kb TSS window (tssRegion = c(−3000, 3000)) and an mm10 transcript database (TxDb generated from mm10.refGene.gtf and confirmed with TxDb.Mmusculus.UCSC.mm10.knownGene^48^).

Genomic feature composition (e.g., promoter, exon, intron, and other categories) was extracted from annotation statistics, and promoter/exon/intron subclasses were collapsed by summing their frequencies within each sample. Feature distributions for WT and KO were combined and visualized as stacked bar plots using ggplot2.

For pathway analysis, peak-associated gene lists for WT and KO were compared using clusterProfiler^39^ compareCluster with KEGG enrichment^49^ and Benjamini–Hochberg multiple-testing correction. Terms with adjusted P < 0.05 that were uniquely enriched in one condition were retained and visualized as dot plots, with color representing −log(adjusted P).

For locus-level visualization, aligned ATAC-seq reads (WT and KO BAM files) were imported using trackViewer^50^ importBam, and gene models were generated from the mm10 TxDb using geneTrack. For each target gene, the plotting region was defined as the gene span extended by 1 kb upstream and 10 kb downstream. WT and KO coverage tracks were displayed together with gene models using viewTracks, applying consistent y-axis limits per gene.

Transcription factor footprinting was performed using TOBIAS.^51^ Tn5 bias–corrected signals were generated with ATACorrect, footprint scores were calculated using FootprintScores, and differential motif binding between WT and KO was assessed with BINDetect using JASPAR2024 CORE non-redundant motifs, the mm10 genome, and the merged consensus peak set. BINDetect output was imported into R, P values were adjusted using the Benjamini–Hochberg method, and motifs were classified as WT_High or KO_High based on direction and significance of binding change. Differential motif activity was visualized as a volcano plot (x-axis: WT–KO binding change; y-axis: −log10 P value) using ggplot2, with significant motifs labeled using ggrepel.

### Anti-PD-1 antibody treatment

For the anti-PD-1 antibody treatment, TC-1 tumor cells were administered into TOX^Ctrl^ and TOX^cKO^ mice. Starting the following day, anti-PD-1 antibody (clone RMP1-14; Bio-X-Cell) or rat IgG2a (clone 2A3; Bio-X-Cell; isotype control) was administered every 3 days for a total of five doses. Mice were sacrificed on day 14 post-tumor cell injection. Lymphocytes isolated from the spleen and tumor were analyzed via flow cytometry.

### Statistical analysis

The statistical significance was analyzed using two-tailed unpaired Student’s *t* tests and two-way analysis of variance with Sidak’s or Tukey’s correction, as appropriate with the Prism 8.0.2 software (GraphPad). Kaplan–Meier survival curves were analyzed using the Mantel–Cox log-rank test with a 95% CI. The Wilcoxon rank-sum test was employed to determine statistical significance in the scRNA-seq dataset. The data are expressed as mean ± standard error of mean (SEM). The difference was considered statistically significant when the *P* value was less than 0.05 (*), 0.01 (**), 0.001 (***), and 0.0001 (****).

## RESULTS

### TOX is preferentially enriched within tumor-infiltrating regulatory T cells across human and murine tumors

To define the distribution of TOX within TI T cell subsets, we profiled immune infiltrates from patients with head and neck squamous cell carcinoma (HNSCC) and non-small-cell lung cancer (NSCLC) using high dimensional mass cytometry (CyTOF) (**Supplementary Table 1**). Major immune lineages were annotated within live CD45^+^ cells, and downstream analyses focused on CD3^+^ T cells (**Extended Data Fig. 1a**). Across patients, CD8^+^ T cells constituted the predominant T cell fraction, whereas conventional CD4^+^ T cells (Tconvs) and Tregs were less abundant (**Extended Data Fig. 1b**). Unsupervised clustering of TI T cells resolved CD8^+^ T cells, Tconvs, and Tregs (**Fig. 1a**). Mapping TOX protein signal onto the same UMAP visualization revealed preferential enrichment of TOX within the Treg compartment, with detectable TOX also present in a subset of CD8^+^ T cells in both cancer types (**Fig. 1a**). Consistent with this pattern, quantitative comparisons across cohorts showed significantly higher TOX expression in Tregs than in CD8^+^ T cells or Tconvs (**Fig. 1b**). TOX^+^ Tregs co-expressed multiple inhibitory receptors and activation-associated molecules, including PD-1, TIM-3, TIGIT, CTLA-4, OX40, and ICOS (**Fig. 1b**). These data indicate that TOX expression is associated with a checkpoint-high, activated TI Treg phenotype in human tumors.

**Fig. 1.**
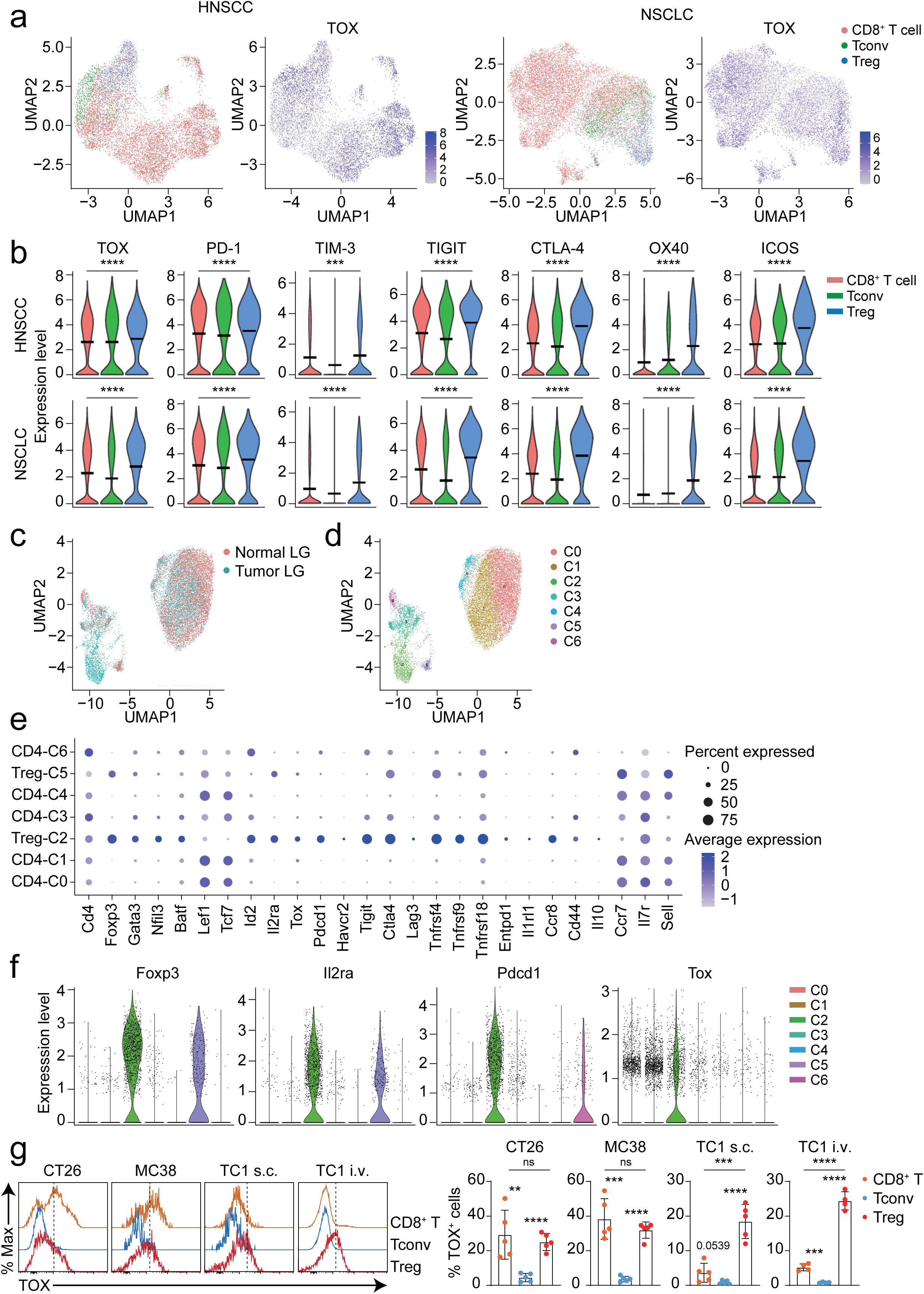
TOX expression is enriched in TI Tregs in both human and murine tumor models. **a**, UMAP visualization of CyTOF analysis from patients with head and neck squamous cell carcinoma (HNSCC) and non-small-cell lung cancer (NSCLC), showing distinct T-cell subsets—CD8⁺ T cells (orange), conventional CD4⁺ T cells (Tconv; green), and regulatory T cells (Treg; blue). The right panels display the scaled expression of TOX across these subsets. **b**, Violin plots depicting the expression of TOX, PD-1, TIM-3, TIGIT, CTLA-4, OX40, and ICOS in CD8^+^ T cells, Tconv, and Treg subsets from HNSCC (top) and NSCLC (bottom), as determined by CyTOF. **c**, UMAP representation of public scRNA-seq data (GSE 152022) of CD4^+^ T cells isolated from normal lung (reddish orange) and tumor lung (teal) tissue, demonstrating distinct immune cell clustering. **d**, UMAP representation showing unsupervised clustering (C0–C6) of CD4^+^ T cells based on single-cell transcriptomic profiles, corresponding to distinct immune subpopulations. **e**, Dot plot summarizing expression of representative marker genes across CD4⁺ T-cell clusters. Dot size indicates the proportion of expressing cells; color intensity represents average expression. Treg clusters (C2 and C5) exhibit high expression of canonical Treg markers, including *Foxp3* and *Il2ra*. **f**, Violin plots illustrating the expression levels of *Foxp3*, *Il2ra*, *Pdcd1*, and *Tox* in each immune cell cluster (C0–C6). **g**, Representative histograms (left) and quantitative analysis (right) of TOX expression in CD8⁺ T cells (orange), Tconvs (blue), and Tregs (red) from CT26, MC38, and TC-1 tumor models. Percentages indicate TOX⁺ cells within each population. Data are representative of three independent experiments (**g**). All bar graphs show mean ± SEM. Statistical significance was determined by two-tailed unpaired Student’s *t*-test (**g**). ns, not significant; **, *P* < 0.01; ***, *P* < 0.001; ****, *P* < 0.0001. s.c., subcutaneous; i.v., intravenous; HNSCC, head and neck squamous cell carcinoma; NSCLC, non-small-cell lung cancer; LG, lung.

To assess whether this enrichment was conserved at the transcriptional level, we analyzed a previously published single-cell RNA sequencing dataset (GSE152022) of murine CD4^+^ T cells obtained from normal lung and TC-1 tumor-bearing lung tissue. UMAP visualization demonstrated distinct cellular distributions between normal and tumor lung compartments (**Fig. 1c**). Unsupervised clustering identified seven CD4^+^ T cell clusters (C0-C6) (**Fig. 1d**), among which clusters C2 and C5 were annotated as Tregs based on high expression of canonical markers such as *Foxp3* and *Il2ra* (**Fig. 1e, f**). *Tox* transcripts were selectively enriched in the tumor-derived Treg cluster relative to the normal lung-derived Treg cluster, with minimal expression across non-Treg CD4^+^ populations (**Fig. 1f**). Within the tumor-derived Treg compartment, *Tox* expression was associated with higher expression of inhibitory receptor transcripts including *Pdcd1*, *Tigit* and *Ctla4*, and activation-associated genes including *Cd44* and selected *TNFR* family members (**Fig. 1e, f**). These data support an association between *Tox* and an activated, checkpoint-associated TI Treg transcriptional state.

We next tested whether preferential TOX expression in TI Tregs was conserved across syngeneic murine tumor models. Flow cytometric analysis of tumor-infiltrating lymphocytes (TILs) from CT26, MC38, and TC-1 tumors confirmed that TOX was significantly enriched in TI Tregs compared with Tconvs, with variable TOX expression in CD8^+^ T cells depending on the tumor model (**Fig. 1g**). In the TC-1 tumor model, the frequency of TOX^+^ cells was significantly higher in TI Tregs than in CD8^+^ T cells (**Fig. 1g**). Moreover, stratification of TI Tregs by PD-1 and TIM-3 co-expression revealed a stepwise increase in TOX across PD-1^−^TIM-3^−^, PD-1^+^TIM-3^−^, and PD-1^+^TIM-3^+^ subsets (**Extended Data Fig. 1c**), linking TOX to progressive acquisition of a checkpoint-high TI Treg phenotype.

Together, these findings establish that TOX is selectively enriched in TI Tregs across human cancers and murine tumor models, and that its expression is tightly linked to a checkpoint-high, activation-associated TI Treg state, prompting us to examine whether TOX functionally contributes to this phenotype.

### Progressive expression of TOX and Treg-associated molecules in TI Tregs during tumor progression

Given its selective enrichment of TOX in TI Tregs, we next examined how TOX expression evolves during tumor progression. TC-1 tumor cells were administered intravenously into wild-type mice to generate lung cancer and cohorts of mice and naïve mice as control were sacrificed at days 7, 14, and 20 for kinetic analysis of TILs in the lung. The frequency of TOX^+^ cells increased progressively in TI Tregs, with a marked rise at day 14 and 20, whereas TOX induction in TI CD8^+^ T cells was modest and delayed (**Fig. 2a**). At day 14 and day 20, TOX^+^ frequencies were higher in TI Tregs than in TI CD8^+^ T cells (**Fig. 2a**). Notably, PD-1 expression in TI Tregs increased in parallel with TOX over the same time course (**Fig. 2a**), consistent with coordinated upregulation of TOX and PD-1 expression during tumor progression.

**Fig. 2.**
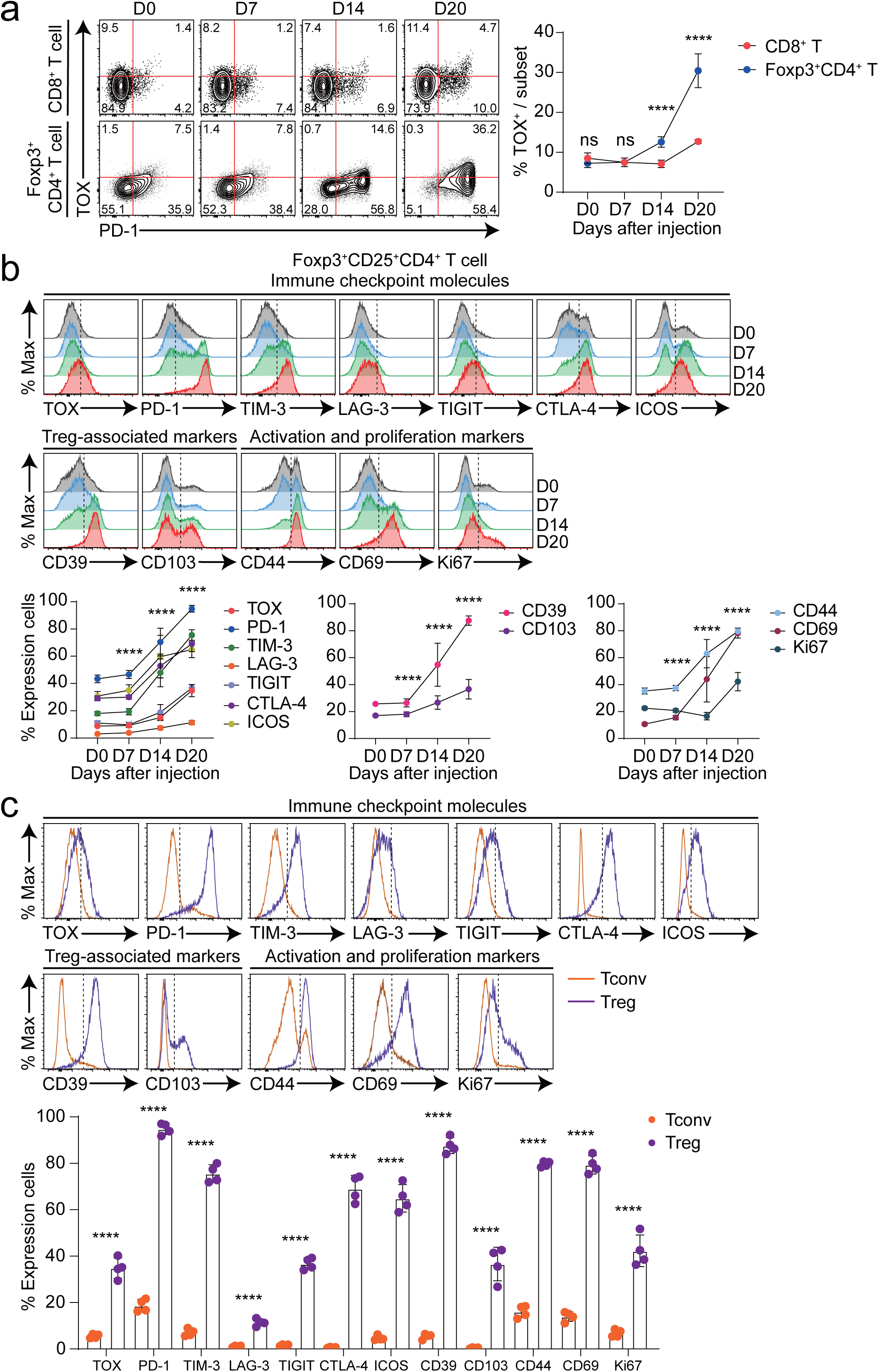
Temporal evolution of TOX and Treg-associated molecules during tumor progression. **a**, Representative flow cytometry contour plots show TOX and PD-1 expression in CD8^+^ T cells and Foxp3^+^CD4^+^ Tregs (left) at days 0, 7, 14, and 20 after TC-1 tumor implantation. Line graph quantifies the increasing frequency of TOX-expressing cells within CD8^+^ T cell (red) and Treg (blue) compartments over time. **b**, Histograms show expression kinetics of TOX, immune checkpoint molecules (PD-1, TIM-3, LAG-3, TIGIT, CTLA-4, ICOS), Treg-associated markers (CD39, CD103), and activation/proliferation markers (CD44, CD69, Ki67) on Tregs at indicated time points post-tumor injection. Corresponding line graphs summarize frequencies of marker-positive Tregs during tumor progression. **c**, Comparative histogram overlays of TOX and Treg-associated markers on TI Tconvs (orange) versus Tregs (purple) at day 20 post-injection. Bar graph below depicts the proportion of marker-positive cells in each subset within the TME. Data were concatenated in each group. Numerical values on plots denote cell population percentages (**a**). Data are representative of two independent experiments. All bar graphs show mean ± SEM. Statistical analyses were performed using two-way ANOVA with Sidak’s *post hoc* test (**a**), Tukey’s *post hoc* test (**b**), and two-tailed unpaired Student’s *t*-test (**c**). ns, not significant; ****, *P* < 0.0001.

We next examined whether increasing TOX expression was accompanied by coordinated changes in other markers associated with TI Tregs. Across time points, TI Tregs showed progressive upregulation of checkpoint molecules - including PD-1, TIM-3, LAG-3, TIGIT, CTLA-4, and ICOS – as well as Treg-associated markers including CD39 and CD103, and activation or proliferation markers including CD44, CD69, and Ki67 (**Fig. 2b**). Marker visualization within TI Tregs further indicated that TOX^+^ cells were enriched among populations expressing these markers (**Extended Data Fig. 1d**).

To place these findings in context, we directly compared TI Tregs and Tconvs within the TME at day 20. TOX expression was significantly higher in TI Tregs than in Tconvs (**Fig. 2c**). A similar pattern was observed for checkpoint molecules, Treg-associated markers, and activation or proliferation markers, all of which were more highly expressed in TI Tregs relative to Tconvs at this timepoint (**Fig. 2c**).

Together, these findings demonstrate that TOX is progressively induced in TI Tregs during tumor progression and is kinetically coupled with the acquisition of a checkpoint-high, effector-like phenotype, supporting its role as an inducible regulator rather than a constitutive feature of TI Treg identity.

### Treg-specific TOX deletion reduces tumor burden and enhances anti-tumor immunity

To directly assess the Treg-intrinsic contribution of TOX to tumor control, we deleted *Tox* selectively in Foxp3^+^ Tregs by crossing *Tox*^fl/fl^ with *Foxp3*^YFP-Cre^ mice to generate Treg-specific TOX-deficient mice (hereafter TOX^cKO^); *Tox*^fl/fl^ littermates served as controls (hereafter TOX^Ctrl^) (**Extended Data Fig. 2a**). Following intravenous challenge with TC-1 tumor cells (**Extended Data Fig. 2b**), TOX^cKO^ mice showed reduced tumor burden and improved survival compared with TOX^Ctrl^ mice (**Fig. 3a, b**). These data support a role for Treg-intrinsic TOX in promoting tumor progression. We next assessed how Treg-specific TOX deletion affects T cell composition in the TME and the spleen. Within the TME, CD8^+^ and CD4^+^ T cell frequencies were significantly increased, and Foxp3^+^ Treg frequency was dramatically reduced in TOX^cKO^ mice relative to TOX^Ctrl^ mice (**Fig. 3c**), demonstrating the improved intratumoral immune balance. In the spleen, however, TOX^cKO^ mice showed reduced frequencies of CD8^+^ and CD4^+^ T cells, with a paradoxical increase in Foxp3^+^ Treg frequency (**Extended Data Fig. 2c**), possibly reflecting homeostatic redistribution of Tregs from the tumor to the periphery or compensatory peripheral Treg expansion. TOX^cKO^ TI Tregs showed reduced expression of immune checkpoint molecules including PD-1, TIM-3, and CTLA-4 and activation-associated markers such as CD44, KLRG1, and Ki67, whereas CD103 expression was increased (**Fig. 3d**), potentially reflecting an altered tissue-residency program in TOX-deficient TI Tregs. *Ex vivo* restimulation with PMA and ionomycin revealed increased production of IFN-γ, TNF-α, and IL-2 by both CD8^+^ and CD4^+^ T cells from TME and spleen of TOX^cKO^ mice (**Fig. 3e** **and Extended Data Fig. 2d**), indicating that Treg-specific TOX deletion permits stronger conventional T cell effector function.

**Fig. 3.**
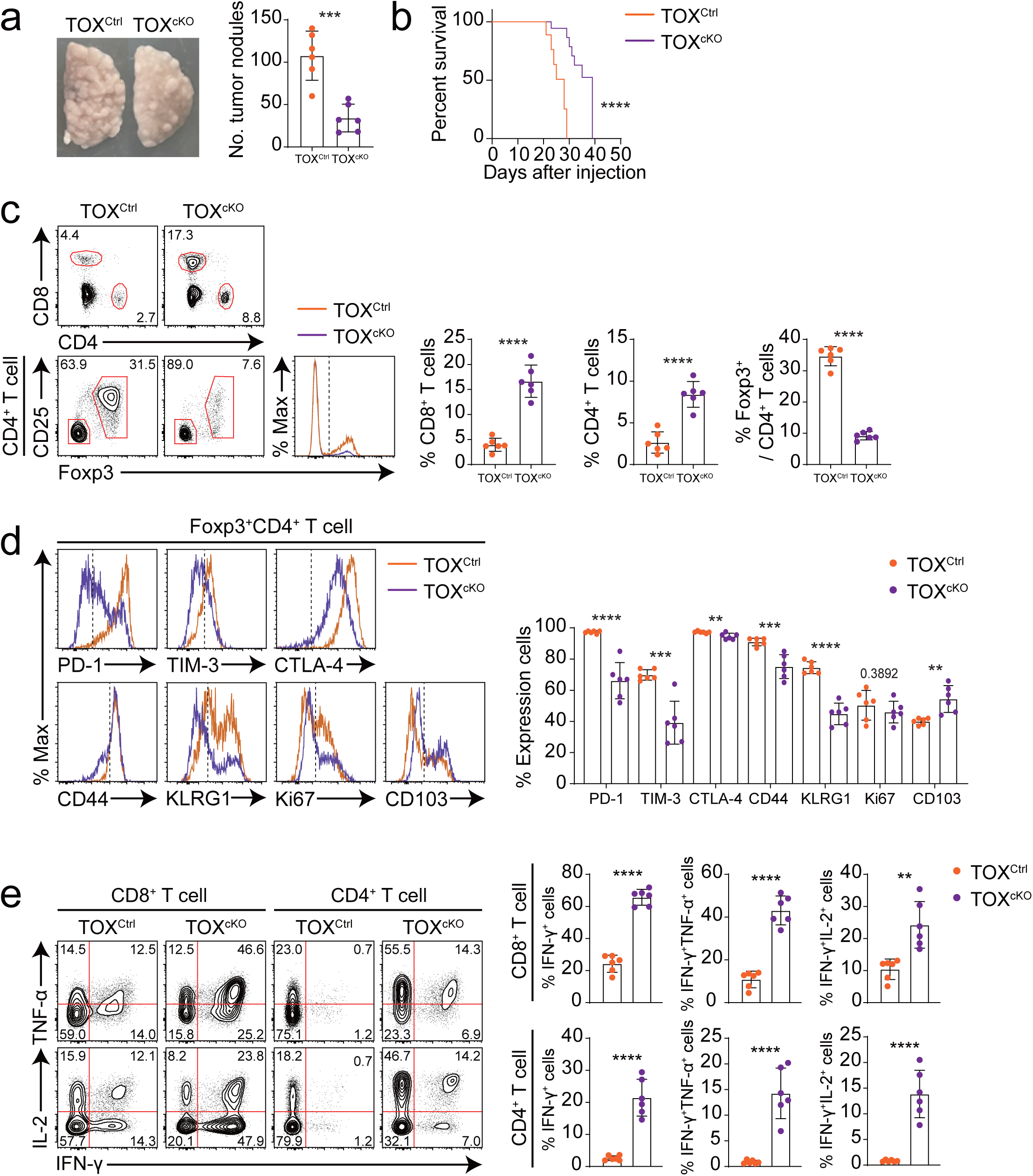
Treg-specific TOX deletion restrains tumor growth and reprograms intratumoral T cell responses. **a**, Representative lung images from TC-1 tumor-bearing TOX^Ctrl^ and TOX^cKO^ mice (left), with quantification of pulmonary tumor nodules per group (*n* = 6/group, right). **b**, Kaplan–Meier survival analysis comparing TOX^Ctrl^ (*n* = 15) and TOX^cKO^ (*n* = 23) cohorts following TC-1 tumor challenge. **c**, Representative flow cytometry analysis of TI CD8^+^ and CD4^+^ T cells and Foxp3^+^ Tregs from TOX^Ctrl^ and TOX^cKO^ mice (*n* = 6 per group). Bar graphs show the proportions of each cell population within tumors between groups. **d**, Representative histograms depict the expression of immune checkpoint molecules (PD-1, TIM-3, and CTLA-4), Treg-associated molecules (KLRG1 and CD103) and the activation/proliferation molecules (CD44 and Ki67) on TI Tregs from TOX^Ctrl^ and TOX^cKO^ (*n* = 6 per group) mice. Bar graphs quantify these markers between groups. **e**, Functional profiling of TI T cells demonstrates effector cytokine production (IFN-γ, TNF-α, IL-2) by both CD8⁺ and CD4⁺ T cells in TOX^Ctrl^ and TOX^cKO^ (*n* = 6 per group) mice. Representative contour plots indicate proportions of cytokine-positive cells. Bar graphs quantify the increase in effector cytokine production between groups. Data were concatenated within each group (**d**). The numbers in the plots indicate the percentage of the population (**c, e**). Data are representative of three independent experiments. All bar graphs show mean ± SEM. Statistical significance was determined by two-tailed unpaired Student’s *t*-test (**a, c–e**) or Mantel–Cox log-rank test for survival with a 95% CI (**b**). **, *P* < 0.01; ***, *P* < 0.001; ****, *P* < 0.0001.

To determine whether Treg-specific TOX deletion similarly impacts tumor control in a distinct tumor context, we examined the B16F10 melanoma lung metastasis model (**Extended Data Fig. 2b**). TOX^cKO^ mice showed reduced pulmonary tumor nodule burden compared with TOX^Ctrl^ mice (**Extended Data Fig. 3a**). In the TME, CD8^+^ T cell frequency increased in TOX^cKO^ mice, while CD4^+^ T cell and Treg frequencies were unchanged (**Extended Data Fig. 3b**). In the spleen, CD8^+^ and CD4^+^ T cell frequencies were comparable between groups, whereas Treg frequency was increased in TOX^cKO^ mice (**Extended Data Fig. 3b**). TOX^cKO^ TI Tregs showed reduced expression of immune checkpoint molecules including PD-1, TIM-3, LAG-3, TIGIT, and CTLA-4 (**Extended Data Fig. 3c**), whereas restimulation assays showed increased effector cytokine production by CD8^+^ and CD4^+^ T cells from TME and spleen in TOX^cKO^ mice (**Extended Data Fig. 3d**). Together, these data indicate that Treg-specific TOX deletion improves tumor control across distinct tumor models, though the magnitude and breadth of TI Treg phenotypic changes vary by tumor type.

Next, to determine whether TOX induction in Tregs is strictly restricted to the TME or extends to other settings of persistent antigen stimulation, we used a murine model of chronic LCMV Cl13 infection. Wild-type mice were infected with Cl13 and TOX expression was tracked over time in Tregs and LCMV-specific H-2D^b^GP276^+^CD8^+^ T cells in the spleen and lung. In both tissues, the frequency of LCMV-specific TOX-expressing CD8^+^ T cells peaked at day 15 and remained elevated at day 32, whereas TOX-expressing Treg frequency peaked at day 15 and declined by day 32 (**Extended Data Fig. 4a, b**), indicating distinct kinetics of TOX regulation in these two populations during chronic infection. We also asked whether Treg-intrinsic TOX shapes T cell responses during chronic infection using TOX^Ctrl^ and TOX^cKO^ mice (**Extended Data Fig. 5a**). TOX^cKO^ mice infected with Cl13 showed reduced splenic Treg frequency and reduced immune checkpoint molecule expression on Tregs compared with TOX^Ctrl^ mice at day 15 (**Extended Data Fig. 5b, c**). Virus-specific CD8^+^ and CD4^+^ T cell effector responses were enhanced in TOX^cKO^ mice across multiple cytokine combinations (**Extended Data Fig. 5d**). These results indicate that Treg-intrinsic TOX also shapes T cell immunity during chronic viral infection, suggesting that TOX-dependent Treg programs are not strictly confined to the TME, albeit potentially through partially overlapping mechanisms.

### TOX is required for TI Treg-mediated suppression of CD8^+^ T cell proliferation and effector function

Given the increased conventional T cell activity in TOX^cKO^ tumors, we next directly tested whether TOX supports the suppressive function of TI Tregs *ex vivo*. TI Tregs were isolated from TC-1 tumor-bearing TOX^Ctrl^ and TOX^cKO^ mice and co-cultured them with CTV-labeled CD8^+^ T cells at graded ratios in the presence of anti-CD3/CD28 stimulation (**Fig. 4a**). TOX^cKO^ TI Tregs showed impaired suppressive capacity across all tested ratios, as reflected by a higher percentage of proliferating CD8^+^ T cells and a greater number of cell divisions compared with cultures containing TOX^Ctrl^ TI Tregs (**Fig. 4b**). Consistent with this, CD8^+^ T cells co-cultured with TOX^cKO^ TI Tregs showed increased IFN-γ production following restimulation relative to those cultured with TOX^Ctrl^ TI Tregs (**Fig. 4c**). Together, these data establish that TOX is cell-intrinsically required for the full suppressive capacity of TI Tregs, directly linking TOX expression to Treg-mediated suppression of CD8^+^ T cell proliferation and effector function in the TME.

**Fig. 4.**
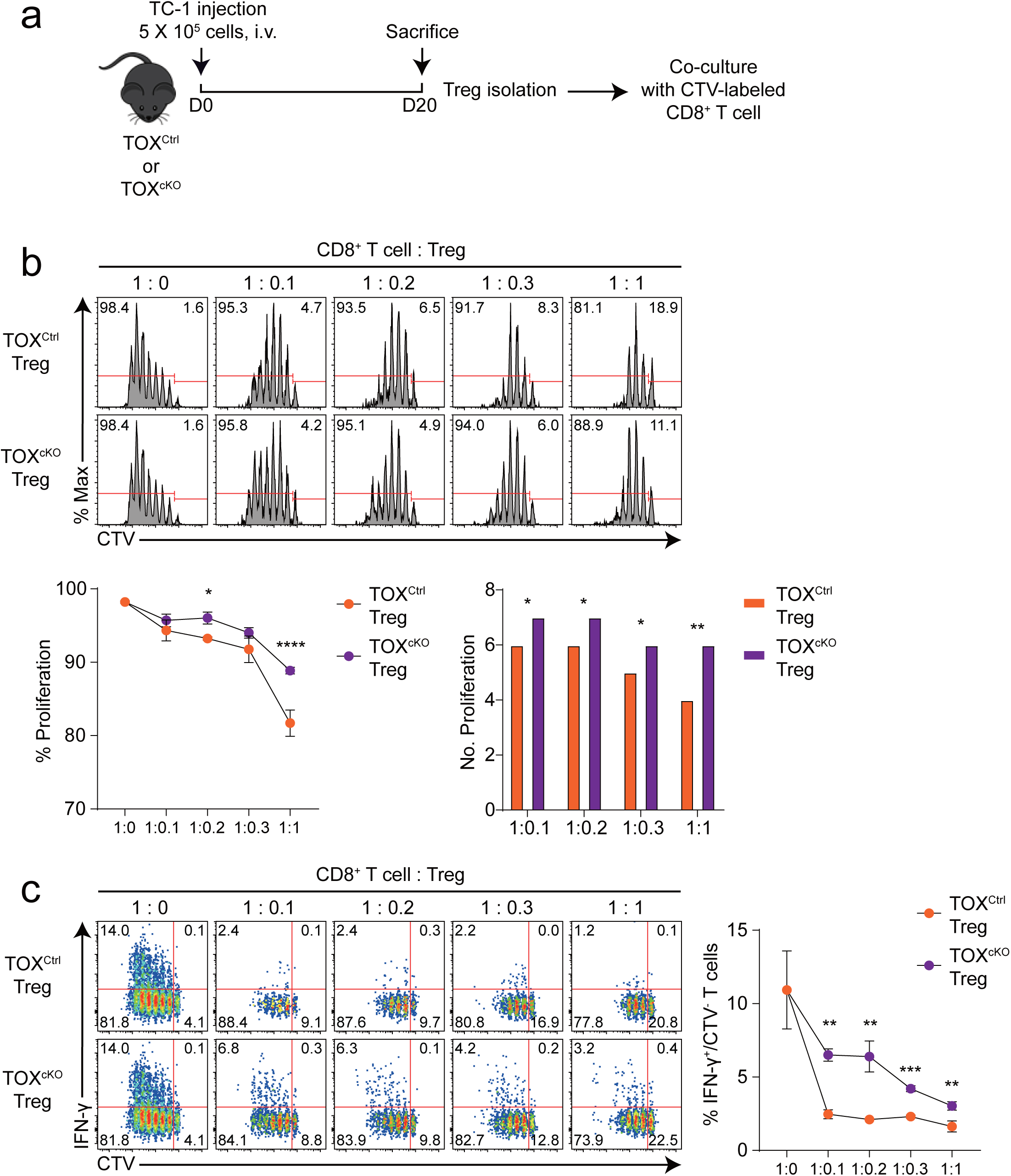
TOX-deficient Tregs display impaired suppressive capacity, leading to enhanced CD8⁺ T-cell proliferation and IFN-γ production. **a**, Experimental design. TOX^Ctrl^ or TOX^cKO^ mice were intravenously inoculated with 5 × 10⁵ TC-1 tumor cells. On day 20, TI Tregs were isolated from each group and co-cultured with CellTrace Violet (CTV)–labeled CD8⁺ T cells at the indicated ratios in the presence of CD3/CD28 Dynabeads for 72 hours to evaluate suppressive capacity. **b**, Flow cytometric analysis of CD8^+^ T cell proliferation, as measured by CTV dilution, in the presence of Tregs from TOX^Ctrl^ or TOX^cKO^ mice. Representative histograms show CD8^+^ T cell proliferation at different CD8^+^ T cell-to-Treg ratios (top). The percentage of proliferating CTV-labeled CD8^+^ T cells is presented in the top left of each histogram. Proliferation curve (bottom left) and bar graph (bottom right) indicate the quantification and number of proliferating CTV-labeled CD8^+^ T cells at different ratios. **c**, Representative flow cytometry plots of IFN-γ production by CD8^+^ T cells co-cultured with TOX^Ctrl^ or TOX^cKO^ Tregs at varying ratios. CTV-labeled CD8^+^ T cells were re-stimulated with soluble anti-CD3 (2μg/ml) and anti-CD28 (2μg/ml) antibodies, along with GolgiPlug/GolgiStop during the final 6 hours of culture. The quantification demonstrates the percentage of IFN-γ^+^CTV^−^CD8^+^ T cells (right). The numbers in the histograms and the plots indicate the percentage of positive population. Data are representative of two independent experiments. All bar graphs show mean ± SEM. Statistical analyses were performed using two-way ANOVA with Sidak’s *post hoc* test (**b**) and two-tailed unpaired Student’s *t*-test (**b, c**). *, *P* < 0.05; **, *P* < 0.01; ***, *P* < 0.001; ****, *P* < 0.0001. i.v., intravenous.

### TOX intrinsically sustains the effector TI Treg state and promotes survival within the TME

Our previous experiments indicated that Treg-specific TOX deletion is associated with impaired TI Treg phenotype and reduced suppressive capacity. However, because TOX^Ctrl^ and TOX^cKO^ mice can differ in tumor burden, these comparisons may confound Treg-intrinsic effects with microenvironmental differences. To control for this variable and directly compare TOX-intact and TOX-deficient Tregs within the same TME, we used female TOX^fl/fl^ × Foxp3^YFP-Cre(+/−)^ mice (hereafter TOX^Cre+/−^). In these mice, random X-chromosome inactivation results in mosaic Cre activity, producing YFP^−^ (TOX^WT^) and YFP^+^ (TOX^KO^) Tregs within the same host and the same TME, thereby enabling direct cell-intrinsic comparisons independent of differences in tumor burden or systemic immune context (**Extended Data Fig. 6a**).

Following TC-1 tumor challenge (**Extended Data Fig. 6b**), YFP^+^CD25^+^ TOX^KO^ Tregs were significantly underrepresented relative to YFP^−^CD25^+^ TOX^WT^ Tregs within the same tumor, resulting in a markedly reduced TOX^KO^-to-TOX^WT^ ratio in the TME (**Fig. 5a**). In this paired setting, TOX^KO^ TI Tregs expressed lower levels of PD-1, TIM-3, and TIGIT and showed reduced CD44 compared with TOX^WT^ TI Tregs (**Fig. 5b**). To extend and confirm this result, TOX^WT^ and TOX^KO^ Tregs were sorted from spleen and tumor and profiled by flow cytometry (**Extended Data Fig. 6c**). TOX^KO^ Tregs showed reduced expression of checkpoint molecules including PD-1, TIM-3, LAG-3, TIGIT, and CTLA-4, as well as reduced CD39, CD44, and Ki67 in both compartments (**Fig. 5c** **and Extended Data Fig. 6d**), demonstrating that TOX loss consistently impairs the acquisition of effector-associated features across tissues.

**Fig. 5.**
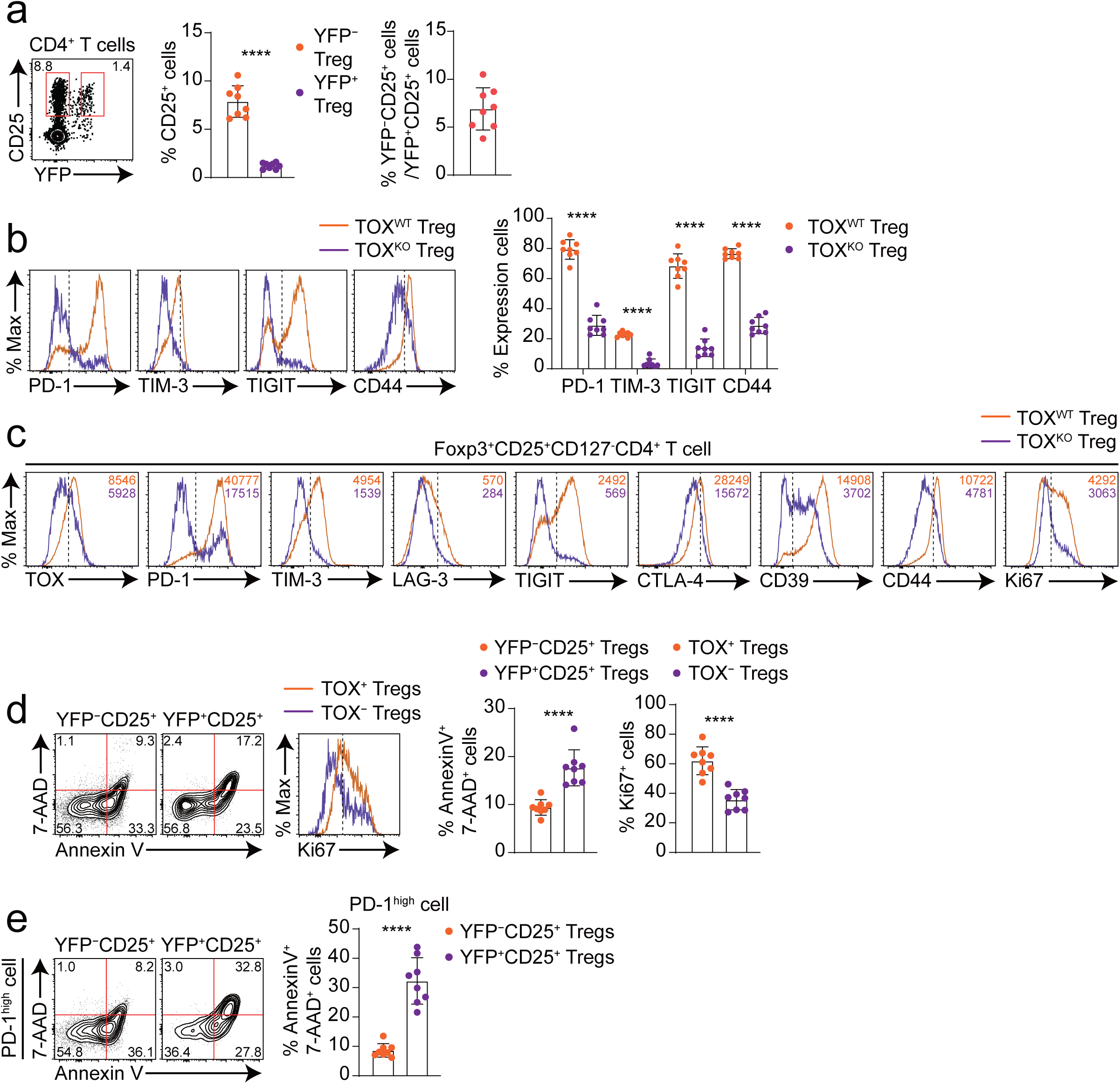
TOX is required for maintaining the effector Treg phenotype and survival in the same TME. **a–b**, Flow cytometric characterization of immune populations and Treg phenotypes in TC-1 tumor-bearing TOX^Cre+/−^ mice. In TOX^Cre+/−^ mice, Foxp3^YFP-Cre+^ cells represent TOX^KO^ Tregs, as Cre-mediated deletion of TOX occurs specifically in Foxp3-expressing cells. Conversely, Foxp3^YFP-Cre−^ cells serve as TOX^WT^ Tregs, retaining normal TOX expression. **a**, Representative contour plot depicts YFP^−^CD25^+^ (TOX^WT^) and YFP^+^CD25^+^ (TOX^KO^) Tregs gated on CD4^+^ T cells from tumor tissue (left). Bar graphs show CD25^+^ Treg frequencies in YFP^−^ and YFP^+^ compartments (middle) and their ratio (right). **b**, Histograms and quantitative analysis of immune checkpoint (PD-1, TIM-3, TIGIT) and activation marker (CD44) expression on Tregs from tumor-infiltrating YFP^−^CD25^+^ (TOX^WT^) and YFP^+^CD25^+^ (TOX^KO^) Treg subsets. **c**, Expanded phenotypic analysis showing the expression of TOX, PD-1, TIM-3, LAG-3, TIGIT, CTLA-4, CD39, CD44, and Ki-67 in sorted TOX^WT^ and TOX^KO^ Tregs from the tumor of TC-1 tumor-bearing TOX^Cre+/−^ mice. **d–e**, Assessment of apoptosis and proliferation in YFP^−^CD25^+^ (TOX^WT^) and YFP^+^CD25^+^ (TOX^KO^) Tregs from TC-1 tumor-bearing TOX^Cre+/−^ mice (*n* = 8). **d**, Flow cytometric analysis (left) and the quantification (right) of apoptotic (Annexin V^+^7-AAD^+^) and proliferating (Ki67^+^) cells among YFP^−^CD25^+^ (or TOX^+^) and YFP^+^CD25^+^ (or TOX^−^) Tregs. **e**, Apoptosis within YFP^−^CD25^+^ and YFP^+^CD25^+^ Tregs gated on PD-1^high^ cells (left) and corresponding quantification of Annexin V^+^7-AAD^+^ frequencies (right). Data were concatenated within each group (**d, e**). Data are representative of two (**a–c**) or three (**d, e**) independent experiments. The numbers in the plots and the histograms indicate the percentage of the population or the MFI (**a, c, d and e**). All bar graphs show mean ± SEM. Statistical significance was determined by two-tailed unpaired Student’s *t*-test. ****, *P* < 0.0001.

We next assessed whether TOX supports TI Treg fitness within this competitive setting. TOX^KO^ TI Tregs showed increased apoptosis and reduced proliferation compared with TOX^WT^ TI Tregs, as indicated by higher Annexin V^+^ 7-AAD^+^ frequencies and lower Ki67^+^ frequencies (**Fig. 5d**). TOX^KO^ Tregs in the spleen also showed increased apoptosis and reduced Ki67 expression compared with TOX^WT^ Tregs (**Extended Data Fig. 6e**), indicating that TOX broadly supports Treg fitness beyond TME. We next examined whether this fitness defect preferentially affected the effector TI Treg compartment. The proportion of PD-1^high^ cells was significantly reduced among TOX^KO^ Tregs in both spleen and tumor (**Extended Data Fig. 6f**), and within the PD-1^high^ gate, TOX^KO^ Tregs showed markedly increased apoptosis compared with TOX^WT^ Tregs in the tumor (**Fig. 5e**) and spleen (**Extended Data Fig. 6g**), indicating that the effector PD-1 ^high^ Treg subset is selectively vulnerable to loss of TOX. Consistent with this, TOX^−^ Tregs displayed a coordinated reduction in effector-associated markers across tissues, spanning checkpoint, suppressive, and activation programs (**Extended Data Fig. 6h**). Together, these data establish that TOX cell-intrinsically sustains the effector-associated TI Treg state and promotes Treg survival within the same TME, ruling out indirect or microenvironmental confounders as primary drivers of the TOX^cKO^ phenotype.

### TOX stabilizes an effector-like, clonally expanded TI Treg state with distinct transcriptional and metabolic programs in TI Tregs

To investigate transcriptional differences between TOX^WT^ and TOX^KO^ Tregs within the same TME, we performed paired single-cell RNA and V(D)J sequencing on TI CD4^+^ T cells from TC-1 tumor-bearing TOX^Cre+/−^ mice. Live CD127^−^CD4^+^ T cells were sorted into YFP^−^CD25^−^, YFP^−^CD25^+^ (TOX^WT^ Tregs), and YFP^+^ (TOX^KO^ Tregs) fractions, labeled with hashtag antibodies, and processed using the 10x Genomics platform (**Extended Data Fig. 7a**). Projection of TI CD4^+^ T cell transcriptomes onto a UMAP identified a *Foxp3* and *Il2ra* expressing Treg compartment within which expression was preferentially enriched (**Fig. 6a**). Unsupervised re-clustering of this Treg compartment resolved seven transcriptional states (**Fig. 6b**), which were assigned to TOX^WT^ or TOX^KO^ origin by hashtag demultiplexing; *Tox* expression was enriched in TOX^WT^ Tregs as expected (**Fig. 6c**).

**Fig. 6.**
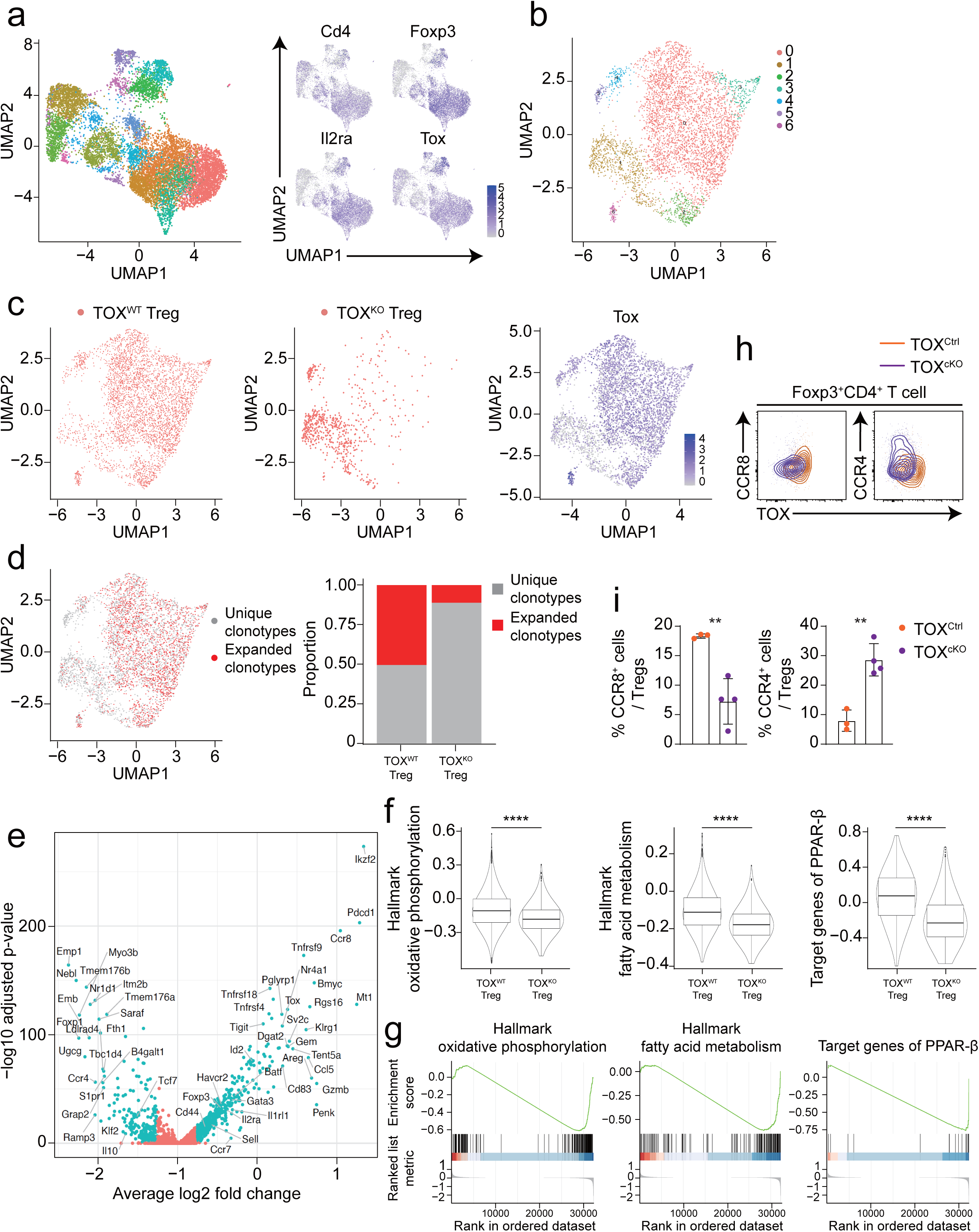
Single-cell transcriptomic and TCR repertoire analyses reveal TOX-dependent transcriptional and metabolic programming of TI Tregs. **a**, UMAP visualization of single-cell transcriptomes from TI CD4⁺ T cells isolated from TC-1 tumor-bearing TOX^Cre+/−^ mice (left). Feature plots depict expression of *Cd4*, *Foxp3*, *Il2ra* and *Tox* across TI CD4^+^ T cell subsets, indicated by colors ranging from gray to blue (right). **b**, UMAP clustering of sorted Tregs from (**a**), revealing seven distinct clusters. **c**, UMAP hashtag demultiplexing distinguishes TOX^WT^ (C0302, left) and TOX^KO^ (C0301, middle) Tregs within tumor-infiltrating populations. Feature plot highlights *Tox* expression (right). **d**, Clonotype analysis visualizes the spatial distribution of unique (grey) and expanded (red) Treg clonotypes (left). Bar graph indicates prevalence of unique versus expanded clonotypes between TOX^WT^ and TOX^KO^ Treg subsets (right). **e**, Volcano plot highlighting differentially expressed genes between TOX^KO^ (left side) and TOX^WT^ (right side) Tregs. **f**, Violin plots illustrate decreased hallmark gene signatures—oxidative phosphorylation, fatty acid metabolism, and PPAR-β target genes—in TOX^KO^ versus TOX^WT^ Tregs. **g**, Gene set enrichment analysis (GSEA) for the hallmark gene sets and target genes of PPAR-β as shown in (**f**), comparing TOX^WT^ versus TOX^KO^ Tregs. **h**, Flow cytometric analysis of CCR8 and CCR4 expression in Foxp3^+^ Tregs from the tumor of TC-1 tumor-bearing TOX^Ctrl^ (*n* = 3) and TOX^cKO^ (*n* = 4) mice, sacrificed on day 20 post-tumor injection. **i**, Quantification of CCR8^+^ and CCR4^+^ Tregs as shown in (**h**). Data were concatenated within each group (**h**). All bar graphs show mean ± SEM. Statistical analyses were performed using Wilcoxon rank sum test (**f**) and two-tailed unpaired Student’s *t*-test (**i**). **, *P* < 0.01; ****, *P* < 0.0001.

V(D)J analysis revealed that TOX^WT^ Tregs harbored a substantially higher proportion of expanded clonotypes, whereas TOX^KO^ Tregs were predominantly composed of unique (non-expanded) clonotypes (**Fig. 6d**), indicating that TOX is associated with clonal expansion of TI Tregs. Differential expression analysis further separated TOX^WT^ and TOX^KO^ programs (**Fig. 6e**): TOX^KO^ Tregs upregulated a stem-like gene program including *Tcf7*, *S1pr1*, and *Klf2*, whereas TOX^WT^ Tregs preferentially expressed an effector-associated program marked by *Foxp3*, *Il2ra*, *Ikzf2*, *Pdcd1*, *Tigit*, and *Gzmb*, together with *Tox* itself (**Fig. 6e**).

Because TI Tregs adapt metabolicaly to the TME, we asked whether TOX status aligned with metabolic gene programs. Gene set variation analysis showed reduced oxidative phosphorylation, fatty acid metabolism, and PPAR-β target gene signatures in TOX^KO^ versus TOX^WT^ Tregs (**Fig. 6f**), and gene set enrichment analysis supported this pattern across the same gene sets (**Fig. 6g**), linking TOX to an effector-associated metabolic state characterized by oxidative and lipid metabolism.

The single-cell data suggested a shift in chemokine receptor usage, with *Ccr8* enriched in TOX^WT^ Tregs and *Ccr4* enriched in TOX^KO^ Tregs (**Fig. 6e** **and Extended Data Fig. 7b**). We validated this prediction by flow cytometry in TC-1 tumor-bearing TOX^Ctrl^ and TOX^cKO^ mice: CCR8^+^ TI Tregs were significantly reduced in TOX^cKO^ tumors, whereas CCR4^+^ TI Tregs were increased (**Fig. 6h, i**), supporting a TOX-dependent bias toward a CCR8-associated effector-like TI Treg phenotype. This is consistent with prior reports identifying CCR8 as a marker of suppressive TI Tregs in human tumors.^9^, ^10^ CCR4^+^ TI Tregs have also been reported to exhibit PD-1^+^TCF1^+^ stem-like properties associated with IL-10 production,^52^ compatible with the CCR4-high, Tcf7-associated program observed in TOX^KO^ TI Tregs.

Because PD-1 signaling has been implicated in TI Treg maintenance and metabolic adaptation, we integrated our dataset with a previously published PD-1^WT^ and PD-1^KO^ TI Treg single-cell dataset (GSE164033) to position TOX deficiency relative to PD-1 deficiency. Integrated clustering separated TOX^KO^ and PD1^KO^ Tregs into distinct clusters, whereas TOX^WT^ and PD-1^WT^ Tregs co-localized within a WT compartment (**Extended Data Fig. 7c, d**). V(D)J analysis supported this separation, with expanded clonotypes enriched in the WT compartment and unique clonotypes predominating in TOX^KO^ and PD-1^KO^ clusters (**Extended Data Fig. 7e–g**). Both TOX^KO^ and PD-1^KO^ clusters shared increased expression of stem-like genes including *Tcf7*, *Klf2*, *S1pr1*, and *Bach2*, whereas the WT effector-like compartment retained higher expression of effector-associated Treg genes including *Tox*, *Tigit*, *Havcr2*, *Tnfrsf4*, *Tnfrsf9*, *Ccr8*, *Batf*, *Id2*, and *Gzmb* (**Extended Data Fig. 7h**). Comparison of differentially expressed genes identified 443 shared DEGs between WT^high^ versus PD-1^KO^ and WT^high^ versus TOX^KO^ Tregs, including *Tigit*, *Ccr8*, *Id2*, and *Ctla4*, together with PD-1^KO^-specific (562 genes; for example, *Ifng*, *Egr4*, and *Cxcr6*) and TOX^KO^-specific (230 genes; for example, *Ikzf4*, *Dusp10*, *Il1rl1*, *Cd80*, *Sell*, and *Tbx21*) gene sets (**Extended Data Fig. 8a**). The total number of DEGs was greater for WT^high^ versus PD-1^KO^ (995 genes) than for WT^high^ versus TOX^KO^ (663 genes) (**Extended Data Fig. 8a**), suggesting that PD-1 deficiency results in a broader transcriptional divergence from the effector TI Treg state. We next visualized these DEG sets across WT^high^, TOX^KO^, and PD-1^KO^ Tregs, which could separate genes into a shared WT^high^ module and additional modules preferentially associated with either PD-1^KO^ or TOX^KO^ (**Extended Data Fig. 8b**). To investigate how these shared and distinct gene programs are reflected at the functional level, we positioned each hallmark pathway by its relative enrichment score along the PD-1 and TOX axes (**Extended Data Fig. 8c**). Pathways linked to the WT^high^ state included angiogenesis and IL2-STAT5 signaling, while PD-1-associated pathways included apoptosis and IL6-JAK-STAT3 signaling, and WNT-β-catenin signaling and TGF-β signaling tracked more closely with the TOX axis (**Extended Data Fig. 8c**).

Together, these single-cell transcriptomic and TCR analyses establish that TOX^WT^ TI Tregs occupy an effector-associated state marked by CCR8 enrichment, higher clonal expansion, and elevated oxidative phosphorylation and fatty acid metabolism gene programs, whereas TOX^KO^ TI Tregs preferentially exhibit a less differentiated, TCF7-associated stem-like transcriptional profile in the TME.

### TOX shapes the chromatin landscape of TI Tregs to enforce AP-1-driven effector programs and suppress TCF7-associated regulatory inputs

To determine whether transcriptional changes in TOX-deficient TI Tregs are accompanied by altered chromatin accessibility, we performed ATAC-seq on FACS-sorted TI Tregs from TC-1 tumor-bearing TOX^Ctrl^ and TOX^cKO^ mice (**Fig. 7a**). Global peak annotation revealed a comparable genomic distribution of accessible regions between groups across annotated features including promoters, introns, and distal intergenic regions (**Fig. 7b**), indicating that TOX loss does not globally restructure chromatin architecture. Nevertheless, given that TOX status was associated with a significant transcriptional difference in TI Tregs (Fig. 6), we asked whether TOX selectively remodels the regulatory landscape of TI Tregs without broadly altering global chromatin architecture.

**Fig. 7.**
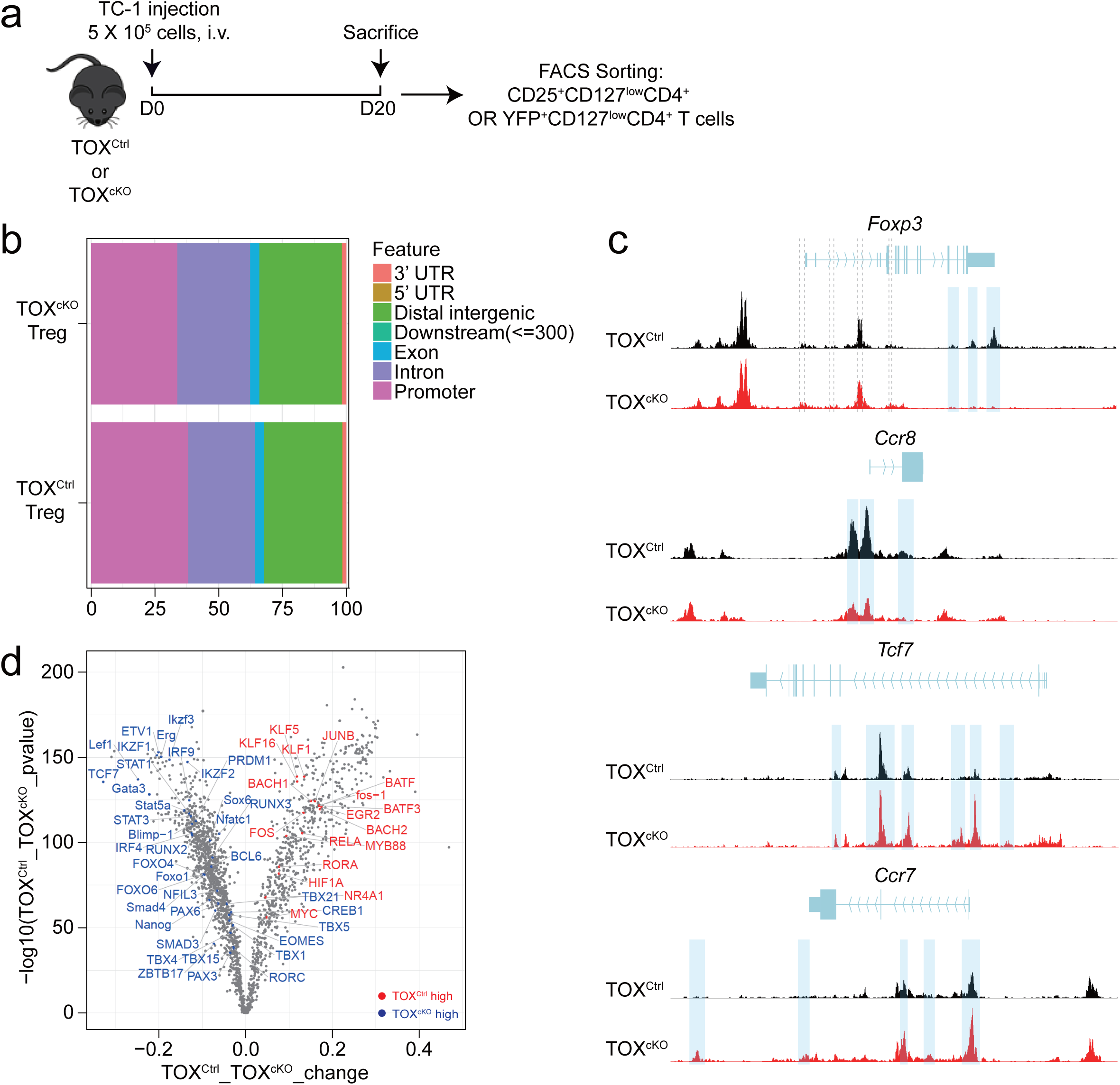
TOX shapes the chromatin landscape of TI Tregs to maintain regulatory and effector programs. **a**, Experimental design. TC-1 tumor-bearing TOX^Ctrl^ and TOX^cKO^ mice were sacrificed on day 20 post-tumor implantation, and TI Tregs (CD25^+^CD127^low^CD4^+^ or YFP^+^CD127^low^CD4^+^ T cells) were sorted by FACS for ATAC-seq analysis. **b**, Genomic distribution of accessible chromatin regions in TOX^Ctrl^ and TOX^cKO^ Tregs, categorized by feature annotation (5’ UTR, 3’ UTR, distal intergenic, downstream, exon, intron, promoter). **c**, Representative ATAC-seq tracks depicting chromatin accessibility at *Foxp3*, *Ccr8*, *Tcf7*, and *Ccr7* loci in TOX^Ctrl^ (black) and TOX^cKO^ (red) Tregs. Blue shaded regions highlight differentially accessible peaks between groups. Dashed vertical lines indicate conserved noncoding sequences (CNS0–3) within the *Foxp3* gene locus. **d**, Volcano plot showing predicted transcription factor binding motif enrichment in sites preferentially accessible in TOX^Ctrl^ high (red, right) or TOX^cKO^ high (blue, left) Tregs.

Consistent with the stem-like transcriptional shift observed in TOX^KO^ TI Tregs by scRNA-seq data (**Fig. 6**), TOX loss was associated with focused accessibility changes at loci linked to TI Treg identity and differentiation state. At the *Foxp3* locus, accessibility at the annotated conserved noncoding sequences (CNS0–3) was comparable between TOX^Ctrl^ and TOX^cKO^ TI Tregs, whereas differential peaks were detected at other regulatory regions within the locus (**Fig. 7c**). In parallel, accessibility at the *Ccr8* locus, a mark of effector-associated TI Tregs, was markedly reduced in TOX^cKO^ TI Tregs (**Fig. 7c**), providing chromatin-level support for the reduced CCR8 protein expression observed by flow cytometry (**Fig. 6h, i**). Conversely, loci associated with stem-like transcriptional programs, including *Tcf7* and *Ccr7*, showed increased accessibility in TOX^cKO^ TI Tregs (**Fig. 7c**). These locus-level changes provide chromatin-level support for a shift away from an effector-associated configuration toward a Tcf7-associated state in the absence of TOX in the TME.

To nominate upstream regulatory inputs linked to these accessibility differences, we performed motif enrichment analysis within accessible regions. Regions preferentially accessible in TOX^Ctrl^ TI Tregs were enriched for *AP-1* family motifs, including *BATF*, *BATF3*, *JUNB*, and *FOS-1*, as well as *NR4A1*, consistent with an activation-driven effector regulatory program (**Fig. 7d**). In contrast, regions preferentially accessible in TOX^cKO^ TI Tregs were enriched for *TCF7*, *LEF1*, and *FOXO1* motifs, together with *STAT* and *IRF* family motifs (**Fig. 7d**), consistent with a shift toward a less differentiated, stem-like regulatory configuration upon TOX loss. KEGG pathway analysis of genes linked to differentially accessible chromatin regions showed that oxidative phosphorylation pathways were enriched among TOX^Ctrl^-associated accessible regions, whereas TOX^cKO^-associated regions pathways were linked to pathways including necroptosis and pyrimidine and lipoic acid metabolism (**Extended Data Fig. 9**), further supporting metabolic and regulatory divergence between TOX^Ctrl^ and TOX^cKO^ TI Tregs.

Collectively, these data establish that TOX enforces an *AP-1*-driven, effector-associated chromatin configuration in TI Tregs, and that its loss redirects the regulatory landscape toward *TCF*-, *LEF1*-, and *FOXO1*-associated programs, providing a chromatin-level mechanism underlying the transcriptional and functional divergence between TOX-sufficient and TOX-deficient TI Tregs in the TME.

### Treg-specific TOX deletion potentiates the efficacy of PD-1 blockade by unleashing intratumoral effector T cell responses

To test whether TOX loss in Tregs can augment PD-1 blockade, TC-1 tumor-bearing TOX^Ctrl^ and TOX^cKO^ mice were treated with isotype control or anti-PD-1 antibody and spleens and tumors were analyzed at day 14 post-tumor challenge (**Fig. 8a**). TOX^cKO^ isotype-treated mice showed significantly reduced tumor burden compared with TOX^Ctrl^ isotype controls, confirming the antitumor effect of Treg-specific TOX deletion (**Fig. 8b**). Anti-PD-1 treatment further reduced tumor burden in TOX^Ctrl^ mice, and the combination of TOX^cKO^ background and anti-PD-1 yielded the lowest tumor burden among all four groups (**Fig. 8b**). Although the incremental reduction conferred by anti-PD-1 over TOX loss alone did not reach statistical significance, possibly reflecting limited statistical power in this experimental setting, the overall pattern was consistent with additive tumor control.

**Fig. 8.**
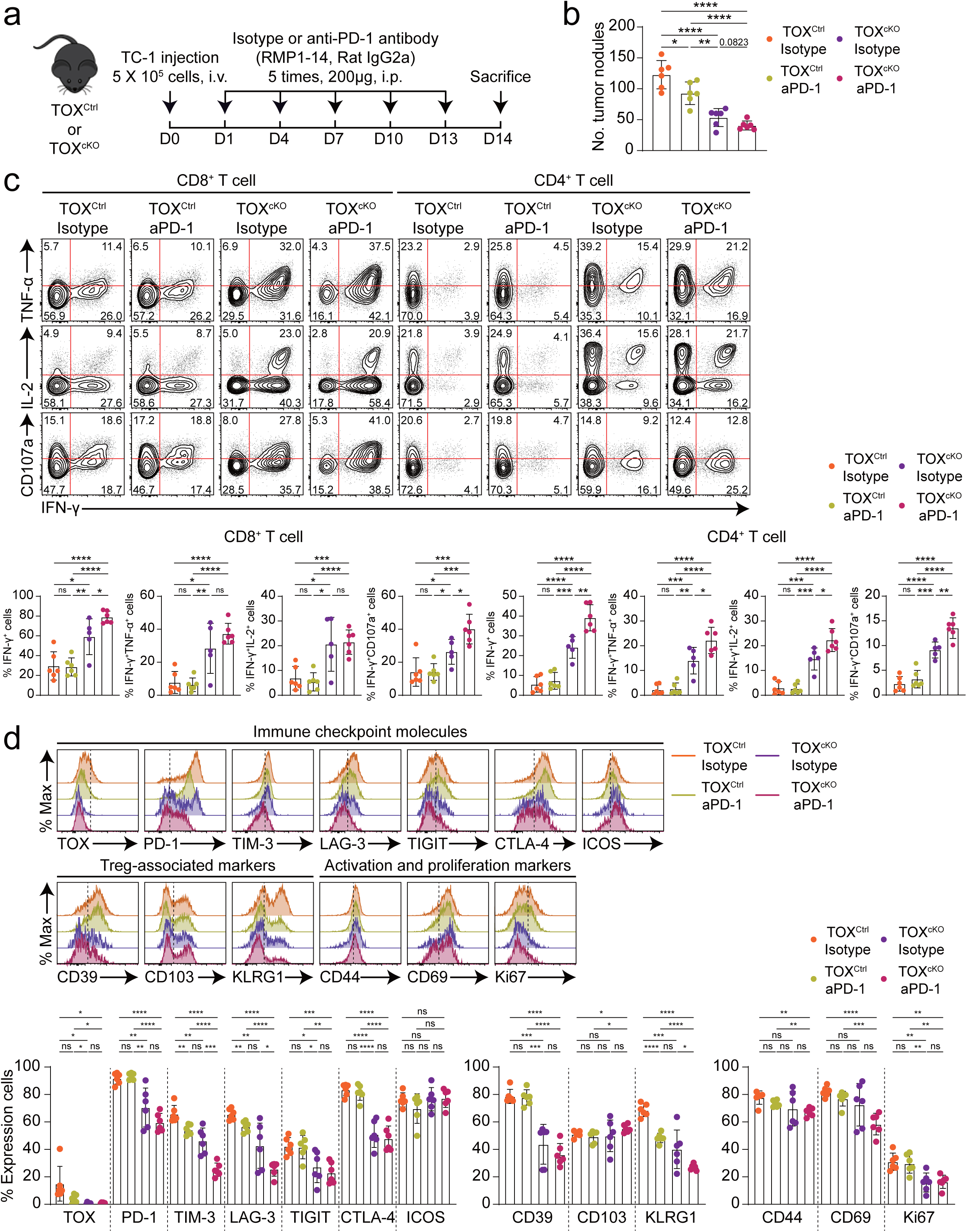
Loss of TOX in Tregs enhances effector T-cell activity and synergizes with PD-1 blockade in the TME. **a**, Experimental design. TC-1 tumor-bearing TOX^Ctrl^ and TOX^cKO^ mice received either isotype control (rat IgG2a, clone 2A3) or anti-PD-1 antibody (RMP1-14, 200 μg, i.p.) on days 1, 4, 7, 10, and 13 following tumor inoculation. Spleen and tumor were harvested at day 14 for immune profiling. Four experimental groups: TOX^Ctrl^ isotype, TOX^Ctrl^ aPD-1, TOX^cKO^ isotype, and TOX^cKO^ aPD-1 (*n* = 6 per group) were analyzed. **b**, Quantification of tumor nodules in TC-1 tumor-bearing mice across four experimental groups: TOX^Ctrl^ isotype, TOX^Ctrl^ aPD-1, TOX^cKO^ isotype, and TOX^cKO^ aPD-1 (*n* = 6 per group). **c**, Flow cytometric analysis of cytokine production (IFN-γ, TNF-α, IL-2) and degranulation marker CD107a in CD8^+^ and CD4^+^ T cells isolated from tumor from each experimental group. Representative contour plots (top) and quantitative bar graphs (bottom) show frequencies of IFN-γ^+^, IFN-γ^+^TNF-α^+^, IFN-γ^+^IL-2^+^, and IFN-γ^+^CD107a^+^ cells among CD8^+^ and CD4^+^ T cells. **d**, Flow cytometric analysis of TOX, immune checkpoint molecules (PD-1, TIM-3, LAG-3, TIGIT, CTLA-4, and ICOS), Treg-associated markers (CD39, CD103, and KLRG1), and activation/proliferation markers (CD44, CD69, and Ki67) in the TI Tregs for all groups. Histograms (top) depict expression profiles across groups, and bar graphs (bottom) quantify the percentage of positive cells for each marker. Data were concatenated for each group (**c, d**). The numbers in the plots indicate the percentage of the population (**c**). All bar graphs show mean ± SEM. Data were analyzed by two-tailed unpaired Student’s *t*-test. ns, not significant; *, *P* < 0.05; **, *P* < 0.01; ***, *P* < 0.001; ****, *P* < 0.0001; i.v., intravenous; i.p., intraperitoneal.

We next assessed conventional T cell function in tumor and spleen by measuring IFN-γ, TNF-α, IL-2, and CD107a expression after *ex vivo* restimulation. In the TME, TOX^cKO^ mice displayed increased frequencies of functional CD8^+^ T cell subsets compared with TOX^Ctrl^ mice, including IFN-γ^+^, IFN-γ^+^TNF-α^+^, IFN-γ^+^IL-2^+^, and IFN-γ^+^CD107a^+^ populations (**Fig. 8c**). Anti-PD-1 treatment further increased these responses most prominently in the TOX^cKO^ background, yielding the highest effector frequencies among the four groups, whereas responses in TOX^Ctrl^ mice remained comparatively restrained (**Fig. 8c**). Similarly, in the spleen, TOX^cKO^ mice a better cytokine response and anti-PD-1 treatment in TOX^cKO^ mice strengthen the effector functions of CD8^+^ T cells (**Extended Data Fig. 10a**). CD4^+^ T cells showed a similar pattern, with the TOX^cKO^ anti-PD-1 group exhibiting the strongest induction of IFN-γ^+^, IFN-γ^+^TNF-α^+^, IFN-γ^+^IL-2^+^, and IFN-γ^+^CD107a^+^ subsets across tumor and spleen (**Fig. 8c** **and Extended Data Fig. 10a**). Together, these data indicate that TOX loss in Tregs is associated with heightened conventional T cell effector activity, and that PD-1 blockade increases the magnitude of these responses most clearly in the TME.

To contextualize these immune changes within the regulatory compartment, we profiled TI Tregs and splenic Tregs across all four groups for checkpoint, Treg-associated, and activation or proliferation-associated markers (**Fig. 8d** **and Extended Data Fig. 10b**). In the tumor, TOX^cKO^ TI Tregs displayed a broad attenuation of the effector-like TI Treg phenotype relative to TOX^Ctrl^, with reduced frequencies of PD-1^+^, TIM-3^+^, LAG-3^+^, TIGIT^+^, and CTLA-4^+^ cells (**Fig. 8d**). This shift extended to additional TI Treg-associated and activation or proliferation markers, including lower CD39, KLRG1, CD44, CD69, and Ki67 in TOX^cKO^ TI Tregs (**Fig. 8d**). Anti-PD-1 treatment imposed marker-specific changes within each genotype, with the most evident effects observed for select inhibitory receptors, rather than uniformly remodeling the TI Treg phenotype (**Fig. 8d**). In the spleen, anti-PD-1 treatment was associated with increased PD-1 expression on Tregs across genotypes, consistent with known compensatory upregulation following PD-1 pathway perturbation (**Extended Data Fig. 10b**). In TOX^Ctrl^ mice, PD-1 blockade was accompanied by upregulation of additional checkpoint molecules including TIM-3, LAG-3, TIGIT, and CTLA-4 on Tregs, suggesting a compensatory activation of alternative inhibitory pathways. By contrast, in TOX^cKO^ mice, anti-PD-1 was associated with more prominent increases in ICOS and CD103, together with higher CD44 and CD69, whereas other checkpoint markers showed smaller or inconsistent changes (**Extended Data Fig. 10b**).

Together, these data demonstrate that TOX-dependent TI Treg programs actively restrain antitumor effector immunity, and that disrupting this program through Treg-specific TOX deletion markedly potentiates the efficacy of PD-1 blockade, providing a mechanistic rationale for targeting TOX-dependent TI Treg circuitry as a strategy to enhance the depth of response to checkpoint immunotherapy.

## DISCUSSION

In this study, we identify TOX as an inducible regulator that is required for TI Tregs to acquire and sustain a checkpoint-high, suppressive state that limits antitumor immunity in the TME. Across human HNSCC and NSCLC, and across multiple syngeneic mouse tumors, TOX was preferentially enriched in TI Tregs relative to conventional T cells, and it rose alongside progressive acquisition of PD-1 and other inhibitory receptors during tumor progression. In contrast, TOX was minimal in Tregs from normal lung tissue in our baseline setting, arguing that TOX is not a constitutive lineage factor but is preferentially associated with the activated TI Treg state that emerges during tumor progression.

Several lines of evidence support a Treg-intrinsic requirement for TOX in maintaining this effector-like TI Treg state. Genetic deletion of TOX in Tregs reduced tumor burden and increased effector function in conventional CD8^+^ and CD4^+^ T cells in the TME. *Ex vivo* suppression assays further showed that TOX-deficient TI Tregs were less able to restrain CD8^+^ T cell proliferation and IFN-γ production. Importantly, by comparing TOX-intact and TOX-deficient Tregs within the same host using mosaic female TOX^Cre+/−^ mice, we could separate Treg-intrinsic effects from differences in tumor burden and microenvironmental intensity. In that paired setting, TOX-deficient Tregs were selectively depleted in tumors and showed reduced expression of checkpoint- and activation-associated features together with increased apoptosis, indicating that TOX supports both the effector-associated phenotype and the persistence of TI Tregs in the same TME (**Fig. 5** **and Supplementary Fig**).

Single-cell transcriptomic and TCR clonotype analyses provided a mechanistic framework for these phenotypes. TOX-intact TI Tregs were enriched for a canonical effector-associated Treg program featuring checkpoint receptors, suppressive effector molecules, and tissue-retention markers, and showed greater clonal expansion. In contrast, TOX-deficient TI Tregs shifted toward a less differentiated program and were dominated by unique clonotypes, consistent with impaired expansion or retention in the tumor. This shift was coupled to reduced oxidative phosphorylation, fatty acid metabolism, and PPAR-β-linked gene programs in TOX-deficient Tregs, connecting TOX-dependent differentiation state to metabolic fitness in the TME. Flow cytometric validation of CCR8 and CCR4 further supported a TOX-dependent bias toward a CCR8-enriched, effector-like TI Treg compartment, whereas TOX loss favored a CCR4-enriched, Tcf7-associated state, consistent with prior reports linking CCR8 to suppressive TI Tregs in human cancers.^9, 10^

Our integrated analysis with published PD-1^WT^ and PD-1^KO^ TI Treg single-cell profiles positions TOX-dependent programming as overlapping with, but not reducible to, PD-1-dependent programming. Both TOX deficiency and PD-1 deficiency were associated with depletion of the effector-like compartment, reduced clonal expansion, and enrichment of a stem-like transcriptional program, consistent with partial convergence on a shared effector Treg module. Notably, PD-1 deficiency was associated with a larger set of differentially expressed genes than TOX deficiency (**Extended Data Fig. 8a**). This difference is compatible with a model in which PD-1 signaling may influence a broader downstream regulatory network that reinforces effector TI Treg programs, whereas TOX controls a more focused, yet critical, set of effector identity and fitness modules, possibly operating in parallel with rather than downstream of PD-1 signaling. Gene-level and pathway-level comparisons indicated separable axes of regulation: angiogenesis and IL-2–STAT5 signaling tracked with the effector-like WT^high^ state, whereas stress and inflammatory signaling pathways preferentially aligned with the PD-1 axis, and WNT–β-catenin and TGF-β signaling aligned more closely with the TOX axis. Together, these data suggest that TOX and PD-1 cooperate to support effector TI Treg biology while controlling non-identical components of the program.

Chromatin accessibility analysis further supports a model in which TOX stabilizes effector-associated regulatory circuitry in TI Tregs. Although global chromatin architecture was comparable between groups, TOX loss was associated with extensive differential accessibility, including reduced accessibility at effector-associated loci such as *Ccr8* and increased accessibility at loci linked to less differentiated states such as *Tcf7* and *Ccr7*. At the *Foxp3* locus, accessibility at CNS elements was largely preserved, whereas differential peaks appeared at other regulatory regions, consistent with maintenance of core lineage identity alongside remodeling of effector-associated features. Motif enrichment within condition-enriched accessible regions nominated distinct transcription factor-associated regulatory states: TOX-intact accessible regions were enriched for *AP-1* family motifs including *BATF*, *BATF3*, *JUNB*, and *FOS*, consistent with activation-coupled effector Treg differentiation. In contrast, TOX-deficient accessible regions were enriched for *TCF7*, *LEF1* motifs and *FOXO1* motifs – the latter being a known regulator of T cell stemness and *TCF7* expression - together with STAT and IRF family motifs, consistent with a shift toward a less differentiated, *FOXO1*-*TCF7*-reinforced regulatory configuration upon TOX loss.

Our findings place TOX-dependent TI Treg programming within a broader framework of lineage-specific TOX function. Recent work has shown that TOX drives CD4^+^ Th1 effector differentiation and IFN-γ production.^53^ Thus, TOX may not impose a single fixed functional output, but may instead operate through a shared axis linking activation-associated differentiation with exit from stem-like states, whose final outcome is shaped by lineage-specific transcriptional and chromatin contexts. Future studies directly comparing TOX-dependent regulatory circuits across T cell lineages will be important for defining this shared regulatory logic and its divergent functional outputs.

In addition to this conceptual implication, our functional data provide a rationale for combinatorial immunotherapy. TOX-deficient Tregs lowered the suppressive barrier in the TME and were associated with heightened effector cytokine and degranulation profiles in conventional T cells. PD-1 blockade produced the strongest functional responses when applied on the background of TOX-deficient Tregs, consistent with a setting in which weakening the effector TI Treg program allows checkpoint blockade to translate into deeper effector activity. The incremental reduction in tumor burden with PD-1 blockade on top of TOX loss was modest in this experiment, which suggests that the degree of synergy may depend on tumor immunogenicity, treatment timing, and the differentiation state of both conventional T cells and Tregs at the time of intervention. Nevertheless, the consistent enhancement of CD8⁺ and CD4⁺ T cell effector function in the TOX^cKO^ anti-PD-1 group supports the concept that selectively disrupting tumor-specific Treg stabilization programs represents a viable strategy to deepen responses to checkpoint immunotherapy.

An important open question is which upstream signals preferentially engage TOX-dependent programs in TI Tregs. The low basal expression of TOX in resting tissue Tregs and its progressive induction during tumor growth and chronic infection point to a requirement for sustained antigen stimulation together with TME-specific inflammatory and metabolic signals. Candidate inputs include persistent TCR signaling, cytokines that engage IL-2–STAT5 pathways, alarmin- and TNF-family signals that promote tissue Treg adaptation, and metabolic stressors such as hypoxia and nutrient competition that are characteristic of tumors. Defining which of these signals is necessary and sufficient to induce TOX, and whether TOX induction requires a specific quality of TCR signaling rather than signal magnitude alone, will be important for understanding when and where TOX-dependent effector Treg programs are formed.

This work has limitations that define clear future directions. First, the TME-derived signals that induce TOX in Tregs remain to be defined. Based on established TOX biology in chronically stimulated T cells, persistent TCR signaling with NFAT-biased activation is a plausible driver, but the contribution of cytokines, hypoxia, metabolites, and antigen-presenting cell programs in the TME should be dissected. Second, whether TOX acts primarily by stabilizing effector-associated enhancer selection, by constraining reversion into resting-associated FOXO1-dominated programs, or by integrating both, will require TOX-centric chromatin-occupancy experiments such as CUT&RUN or ChIP-seq, together with perturbations of candidate upstream regulators. Third, given that high PD-1 can mark distinct functional states in Tregs depending on context, careful temporal and spatial stratification – including analysis of paired pre- and post-treatment tumor biopsies from patients receiving PD-1 blockade – will be important when translating TOX-aligned TI Treg signatures to patient settings and for identifying tumors most likely to benefit from strategies targeting TOX-dependent Treg programs.

In sum, our results identify TOX as an inducible regulator preferentially engaged in TI Tregs that stabilizes an effector-associated suppressive state through coordinated transcriptional and chromatin remodeling with coupled metabolic fitness. This provides a mechanistic explanation for the loss of TI Treg persistence and suppressive function upon TOX deletion in the TME, and establishes a new principle of tumor-specific Treg adaptation driven by inducible transcriptional programming. These findings support therapeutic strategies that selectively disrupt TOX-dependent TI Treg stabilization to enhance the efficacy of checkpoint-based immunotherapy.

## ACKNOWLEDGEMENTS

This work was supported by the National Research Foundation of Korea (NRF) grant funded by the Korea government (MSIT) (RS-2024-00392705, RS-2025-18362970 to S.-J.H.; RS-2024-00411768 to H.R.K.). This work is also supported by the Technology development Program funded by the Ministry of SMEs and Startups (MSS, Korea) (RS-2023-002732224) and by Korea Basic Science Institute (National Research Facilities and Equipment Center) grant funded by the Ministry of Education (RS-2025-02402969). The work was supported in part by Brain Korea 21 (BK21) FOUR program. Supplementary figures were created by BioRender.com. The funders had no role in the study design, data collection and analysis, decision to publish or preparation of the manuscript.

## AUTHOR CONTRIBUTIONS

S.P. conceived the study, established the experimental models, designed and performed the experiments, analyzed the FACS and CyTOF data, wrote and edited the manuscript. D.J.P. assisted with experiments. M.J.K. provided conceptual advice for this study. G.K., R.Z., and S.K.-S. supported CyTOF experiments. G.M.K. prepared human tumor samples. H.R.K. provided human tumor samples. K.K. performed scRNA-seq, scV(D)J-seq, and ATAC-seq data analysis and edited the manuscript. S.-J.H. supervised the overall study and manuscript preparation. J.J., S.K.-S., H.R.K. and S.-J.H. supervised the study and provided guidance on manuscript development. All authors discussed the results, reviewed the manuscript and approved the final version.

---

**Extended Data Fig. 1.**
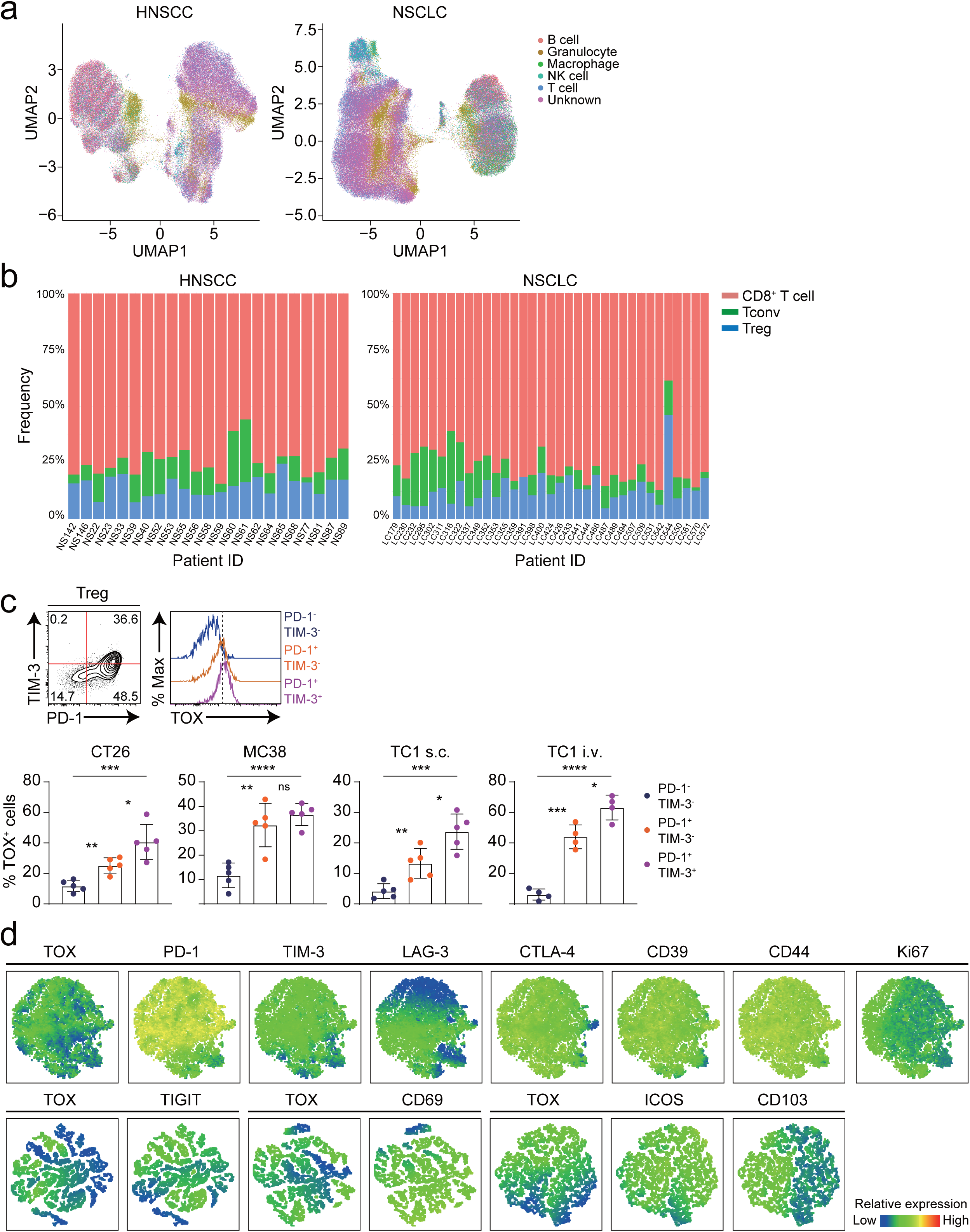
Elevated TOX and Treg-associated molecule expression in TI Tregs (related to Fig. 1 and 2). **a**, UMAP representation of CyTOF data from head and neck squamous cell carcinoma (HNSCC) and non-small cell lung carcinoma (NSCLC) samples following CD45^+^ immune cell isolation. Distinct immune populations were identified using canonical markers: B cells (CD3^−^CD19^+^), granulocytes (CD66b^+^), macrophages (CD3^−^CD68^+^), natural killer (NK) cells (CD3^−^CD56^+^), and T cells (CD3^+^). **b**, Stacked bar plots showing the relative frequencies of CD8⁺ T cells (reddish orange), conventional CD4⁺ T cells (Tconv; green), and regulatory T cells (Treg; blue) across individual HNSCC and NSCLC patients. **c**, Representative flow cytometry analysis of TOX expression within Treg subsets categorized by PD-1 and TIM-3 expression: PD-1^−^TIM^−^ (navy), PD-1^+^TIM-3^−^ (orange), and PD-1^+^TIM-3^+^ (purple) across three tumor models (CT26, MC38, TC-1). Bar graphs represent the proportion of TOX^+^ cells within each Treg subset (bottom). **d**, *t*-SNE visualization depicting relative expression patterns of TOX, immune checkpoint molecules (PD-1, TIM-3, LAG-3, TIGIT, CTLA-4, ICOS), Treg-associated markers (CD39, CD103), and activation/proliferation markers (CD44, CD69, Ki67) in TI Tregs. Data were concatenated within each group (**d**). The numbers in the plots indicate the percentage of the population. Data are representative of two (**d**) or three independent experiments (**c**). All bar graphs show mean ± SEM. Statistical analyses were performed using two-tailed unpaired Student’s *t*-test (**c**). ns, not significant; *, *P* < 0.05; **, *P* < 0.01; ***, *P* < 0.001; ****, *P* < 0.0001. s.c., subcutaneous; i.v., intravenous.

**Extended Data Fig. 2.**
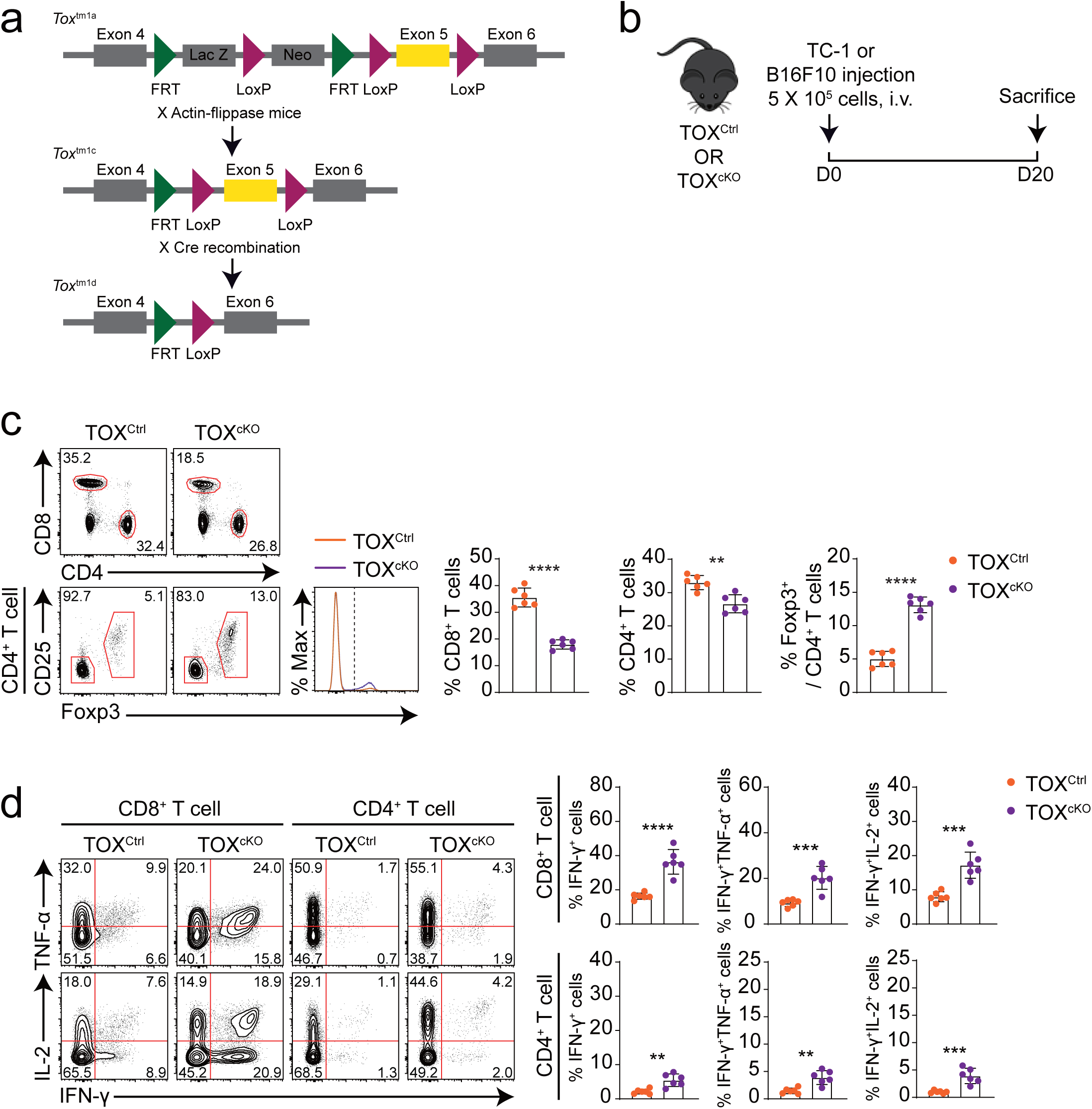
Treg-selective ablation of TOX modulates splenic T cell phenotypes (related to Fig. 3). **a**, Schematic representation of the conditional *Tox* deletion strategy in Foxp3-expressing regulatory T cells. C57BL/6N *Tox*^tm1a^ mice were crossed with *Actin*-Flippase mice to excise the FRT-flanked neomycin resistance cassette, generating *Tox*^tm1c^ (TOX^Ctrl^) mice. *Tox*^tm1c^ mice were subsequently bred with *Foxp3*^YFP-Cre^ reporter mice to produce *Tox*^tm1d^ (TOX^cKO^) mice with Treg-specific deletion of exon 5. **b**, Experimental scheme. TOX^Ctrl^ or TOX^cKO^ mice were intravenously injected with 5 × 10⁵ TC-1 or B16F10 melanoma cells and sacrificed on day 20 for analysis. **c**, Representative flow cytometry plots (left) and quantification (right) of CD8^+^ T cells, CD4^+^ T cells, and Foxp3^+^ regulatory T cells in the spleen of TOX^Ctrl^ and TOX^cKO^ mice bearing TC-1 tumors (*n* = 6 mice per group). **d**, Flow cytometric assessment of cytokine production (IFN-γ, TNF-α, IL-2) by CD8^+^ and CD4^+^ splenic T cells from TOX^Ctrl^ and TOX^cKO^ (*n* = 6 per group), with representative plots and quantified effector cytokine frequencies indicated. The numbers in the plots indicate the percentage of the population (**c, d**). Data are representative of three independent experiments. All bar graphs show mean ± SEM. Data were analyzed by two-tailed unpaired Student’s *t*-test (**c, d**). **, *P* < 0.01; ***, *P* < 0.001; ****, *P* < 0.0001.

**Extended Data Fig. 3.**
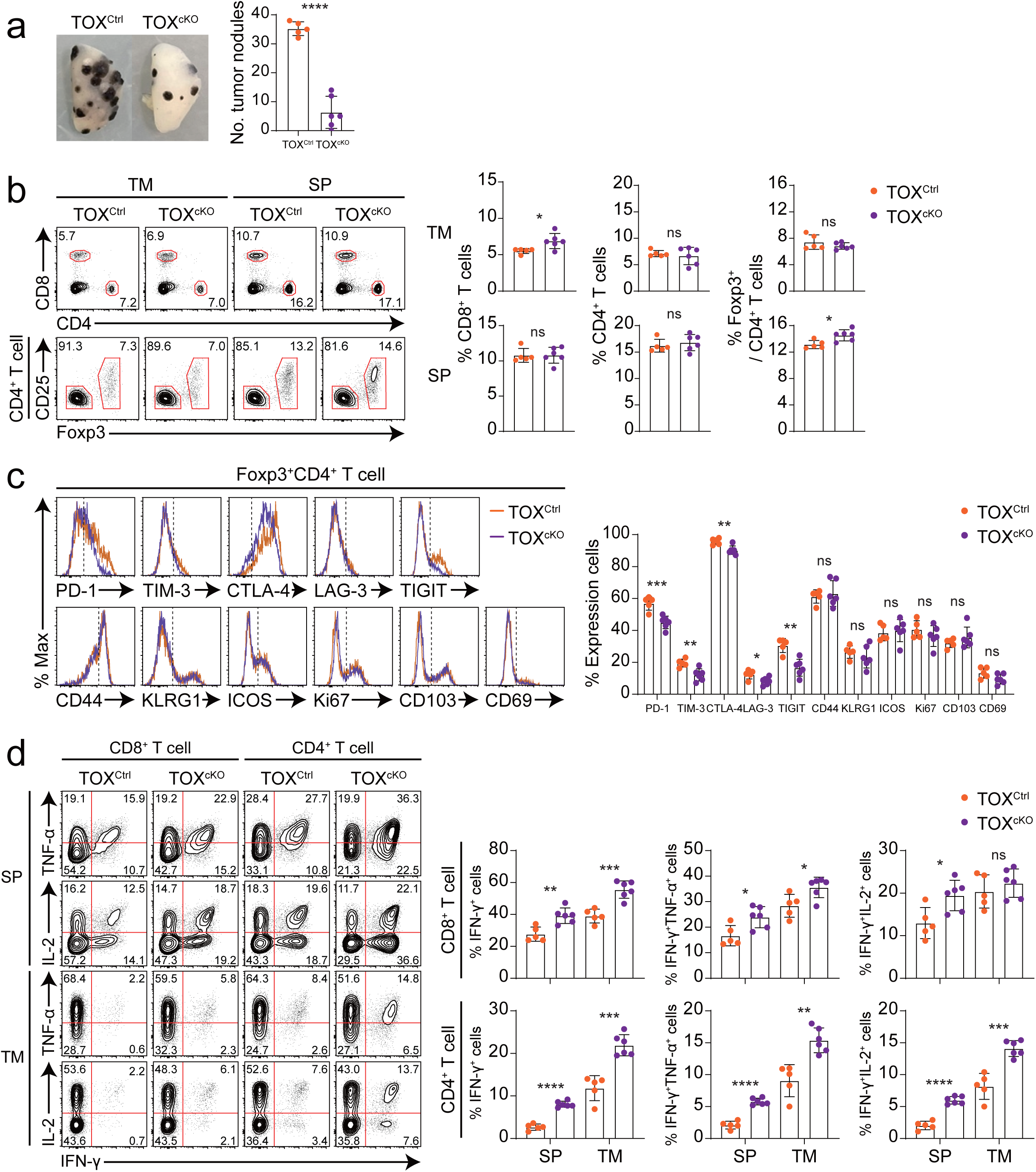
Treg-specific TOX ablation attenuates B16F10 melanoma lung metastasis and disrupts TI Treg phenotype (related to Fig. 3). **a**, Representative images of lungs from TOX^Ctrl^ (*n* = 5) and TOX^cKO^ mice (*n* = 6) following B16F10 tumor challenge, illustrating tumor nodule burden (left). Quantitative assessment of pulmonary tumor nodules at day 20 post-injection is presented (right). **b**, Flow cytometric analysis of CD8⁺, CD4⁺, and Foxp3⁺ T cells in spleen (SP) and tumor (TM) from B16F10-bearing TOX^Ctrl^ and TOX^cKO^ mice as shown in (**a**). Bar graphs indicate population frequencies in each tissue. **c**, Representative histograms (left) and quantification (right) of immune checkpoint molecules (PD-1, TIM-3, LAG-3, TIGIT, CTLA-4, ICOS), Treg-associated markers (CD103, KLRG1), and activation/proliferation markers (CD44, CD69, Ki67) on TI Foxp3^+^ Treg from TOX^Ctrl^ and TOX^cKO^ mice. **d**, Quantification of cytokine production (IFN-γ, TNF-α, IL-2) by CD8^+^ and CD4^+^ T cell subsets in the spleen and tumor from TOX^Ctrl^ and TOX^cKO^ mice. Flow cytometry plots show the percentage of IFN-γ, TNF-α, and IL-2 producing cells (left); bar graphs display the quantified data (right). Data were concatenated in each group (**b–d**). Data are representative of two independent experiments. All bar graphs show mean ± SEM. Data were analyzed by two-tailed unpaired Student’s *t*-test. ns, not significant; *, *P* < 0.05; **, *P* < 0.01; ***, *P* < 0.001; ****, *P* < 0.0001; SP, spleen; TM, tumor.

**Extended Data Fig. 4.**
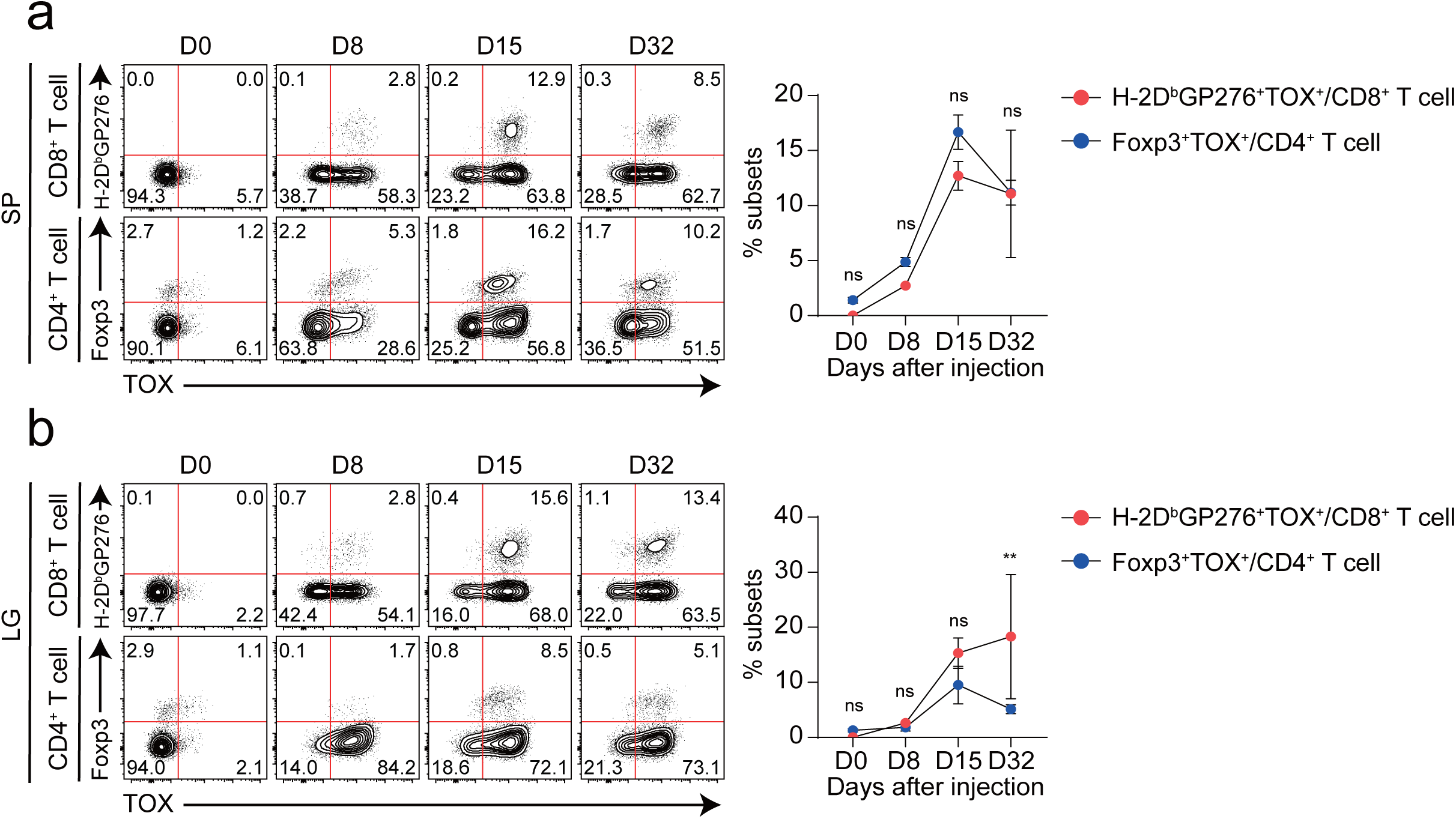
Temporal dynamics of TOX in antigen-specific CD8^+^ T cells and Tregs during chronic LCMV infection. **a**–**b**, Flow cytometric analysis of TOX expression in H-2D^b^GP276^+^CD8^+^ T cells and Foxp3^+^ Tregs from the spleen (**a**) and lung (**b**) at days 0, 8, 15, and 32 post-infection. Representative flow cytometry plots (left) show the percentage of TOX^+^ cells in both CD8^+^ T cells and Tregs. Line graphs (right) quantify the changes in TOX expression over time. Statistical significance was assessed with two-way analysis of variance (ANOVA) with Sidak’s *post hoc* test. ns, not significant; **, *P* < 0.01. i.v., intravenous; SP, spleen; LG, lung.

**Extended Data Fig. 5.**
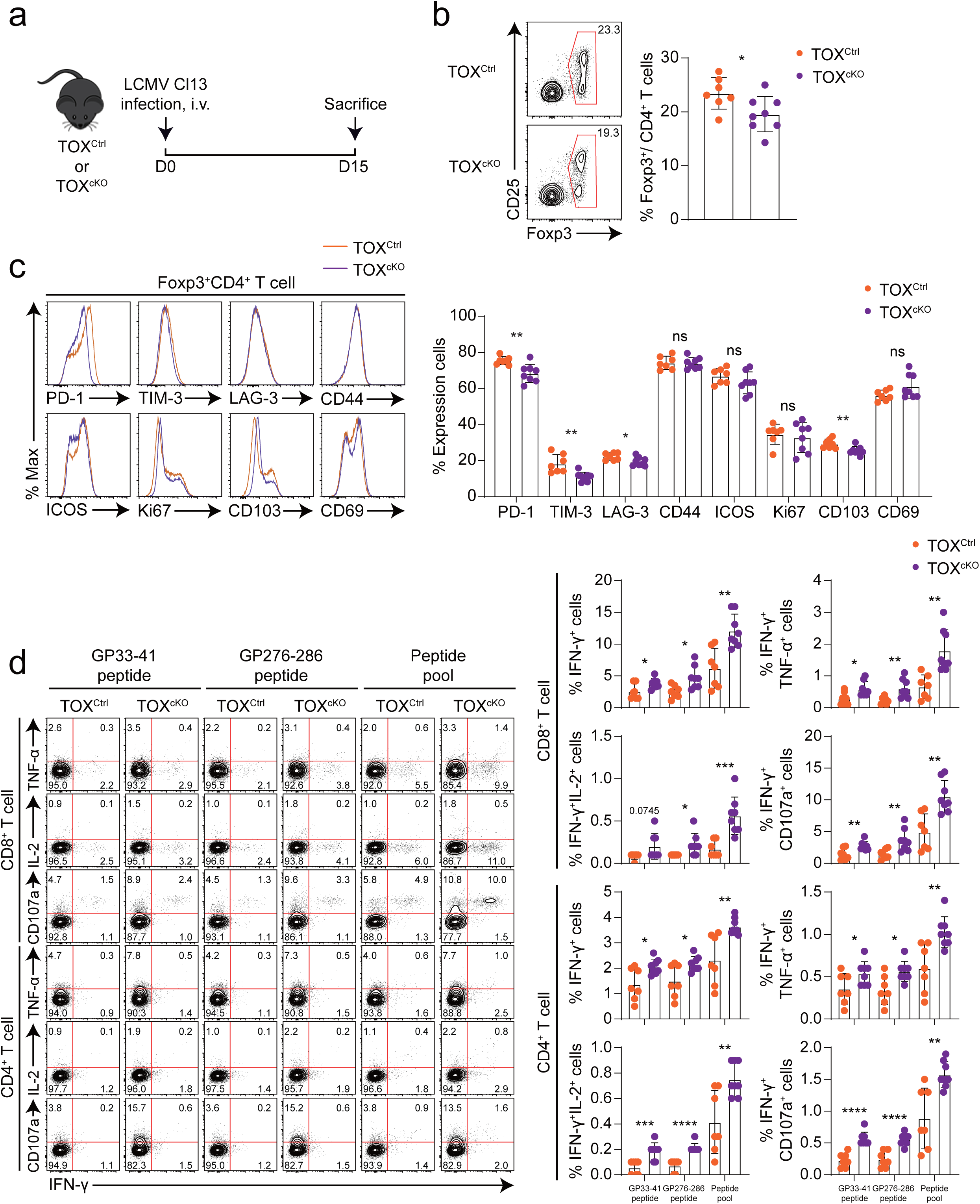
TOX expression in Tregs and its impact on effector T cell responses during chronic viral infection. **a**, Experimental scheme. TOX^Ctrl^ and TOX^cKO^ mice were intravenously infected with 2 × 10^6^ p.f.u. of LCMV Cl13, and spleens collected on day 15 post-infection for immunophenotyping of Tregs and effector T cells. **b**, Representative flow cytometry plots (left) and quantitative analysis (right) displaying the frequency of Foxp3^+^ Tregs in spleens of TOX^Ctrl^ and TOX^cKO^ mice. **c**, Flow cytometric analysis of immune checkpoint molecules (PD-1, TIM-3, LAG-3, and ICOS), and Treg-associated/activation markers (CD103, CD44, CD69, and Ki67) in Foxp3^+^ Tregs from the spleen of TOX^Ctrl^ and TOX^cKO^ mice. Histograms (left) depict the expression profiles, and bar graphs (right) quantify the expression of each marker across groups. **d**, Analysis of cytokine production in CD8^+^ and CD4^+^ T cells from spleens following ex vivo re-stimulation with LCMV-derived peptides (GP33–41, GP276–286, GP66–80, GP70–77, GP92–101, GP118–125). Flow cytometric plots (left) and corresponding bar graphs (right) depict frequencies of IFN-γ^+^, IFN-γ^+^TNF-α^+^, IFN-γ^+^IL-2^+^, and IFN-γ^+^CD107a^+^ cells within CD8^+^ and CD4^+^ T cell subsets. Data were concatenated within each group. The numbers in the plots indicate the percentage of the population (**b, d**). Data are representative of two independent experiments. All bar graphs show mean ± SEM. Data were analyzed by two-tailed unpaired Student’s *t*-test. ns, not significant; *, *P* < 0.05; **, *P* < 0.01; ***, *P* < 0.001; ****, *P* < 0.0001. SP, spleen.

**Extended Data Fig. 6.**
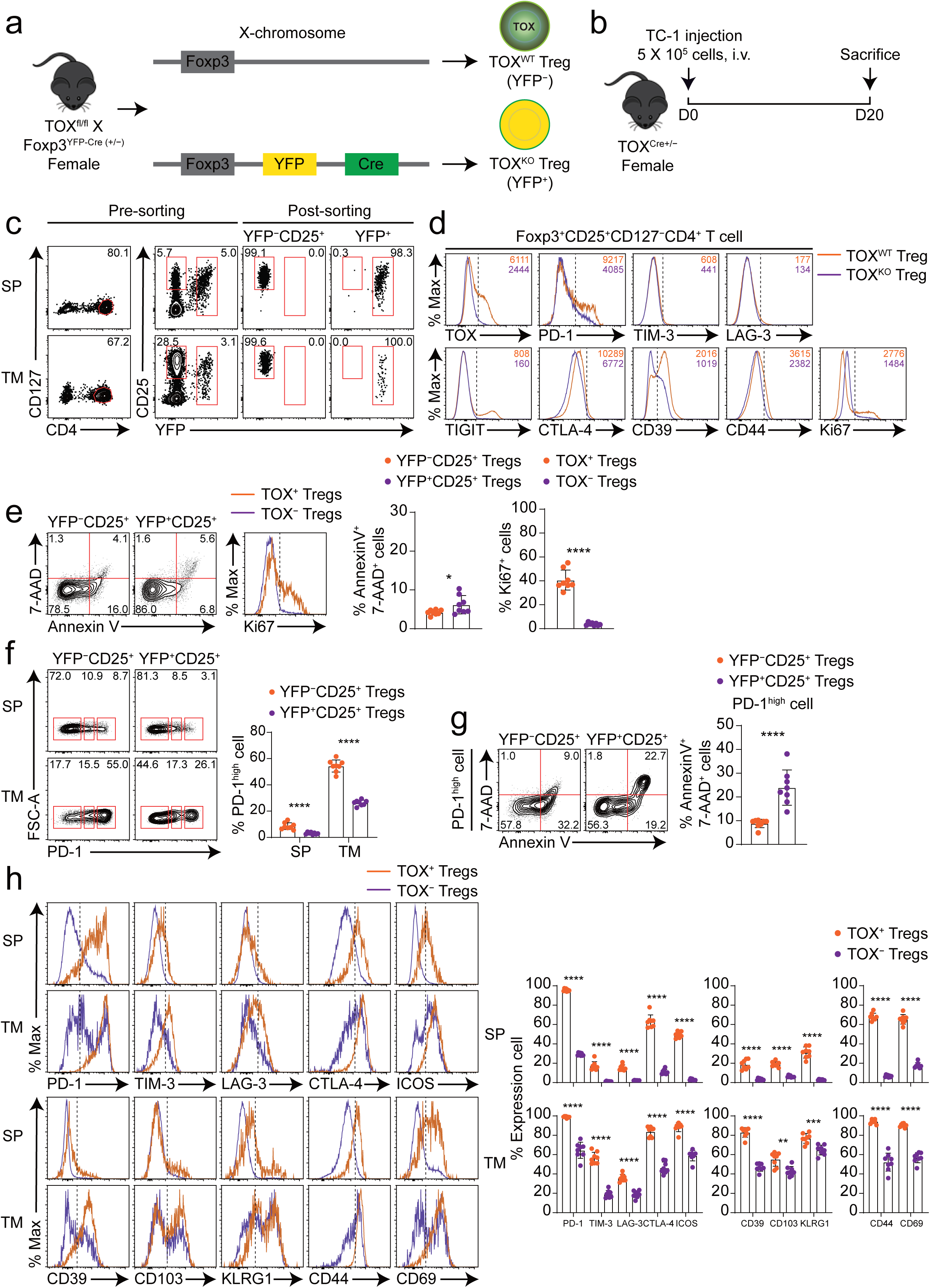
TOX deficiency reshapes TI Treg phenotype (related to Fig. 5). **a**, Schematic representation of the genetic strategy to generate TOX^WT^ (YFP^−^) and TOX^KO^ (YFP^+^) Tregs within the same host. Female TOX^fl/fl^ X Foxp3^YFP-Cre^ ^(+/−)^ mice (TOX^Cre+/−^ mice) harbor both *Tox*-intact and *Tox*-deficient Tregs due to random X-chromosome inactivation in Foxp3-expressing cells. **b**, Experimental scheme. female TOX^Cre+/−^ mice (*n* = 8) were intravenously injected with 5 × 10^5^ TC-1 tumor cells on day 0 and analyzed on day 20 post-injection. **c**, Representative flow cytometry plots showing the gating strategy for fluorescence-activated cell sorting (FACS) of YFP^−^CD25^+^ (TOX^WT^) and YFP^+^ (TOX^KO^) Tregs from pooled spleens and tumors of TC-1 tumor-bearing TOX^Cre+/−^ mice (*n* = 19). Pre-sorting (left) and post-sorting (right) purity are shown. **d**, Representative histograms showing the expression of TOX, immune checkpoint molecules, Treg-associated markers, and activation/proliferation markers on splenic TOX^WT^ and TOX^KO^ Tregs from TC-1 tumor-bearing TOX^Cre+/−^ mice. **e**, Apoptosis and proliferation assessment in YFP^−^CD25^+^ and YFP^+^CD25^+^ Tregs from spleen of TC-1 tumor-bearing TOX^Cre+/−^ mice (*n* = 8). Flow cytometry plots and histograms depict Annexin V^+^7-AAD^+^ apoptotic cells and Ki67^+^ proliferating cells, with bar graphs quantifying frequencies. **f**, Analysis of PD-1^high^ cells in the YFP^−^CD25^+^ and YFP^+^CD25^+^ Tregs from the spleen and tumor of TC-1 tumor-bearing TOX^Cre+/−^ mice (*n* = 8) as shown in Fig. 5e. Flow cytometry plots (left) and bar graphs (right) quantify the frequency of PD-1^high^ cells in these Treg populations. **g**, Representative flow cytometry plots (left) and quantification (right) of apoptotic (Annexin V^+^7-AAD^+^) cells within the PD-1^high^ gate of splenic YFP^−^CD25^+^ and YFP^+^CD25^+^ Tregs from TC-1 tumor-bearing TOX^Cre+/−^ mice (*n* = 8). **h**, Expression of Treg-associated molecules in TOX^+^ and TOX^−^ Tregs from spleen and tumor of TC-1 tumor-bearing TOX^Cre+/−^ mice, illustrated by histograms (left) and quantified expression levels (right). Data were concatenated in each group (**e–h**). Data are representative of two (**e–h**) or three (**c, d**) independent experiments. The numbers in the plots and the histograms indicate the percentage of the population or the MFI (**c–g**). All bar graphs show mean ± SEM. Data were analyzed by two-tailed unpaired Student’s *t*-test. *, *P* < 0.05; ****, *P* < 0.0001. i.v., intravenous; SP, spleen; TM, tumor.

**Extended Data Fig. 7.**
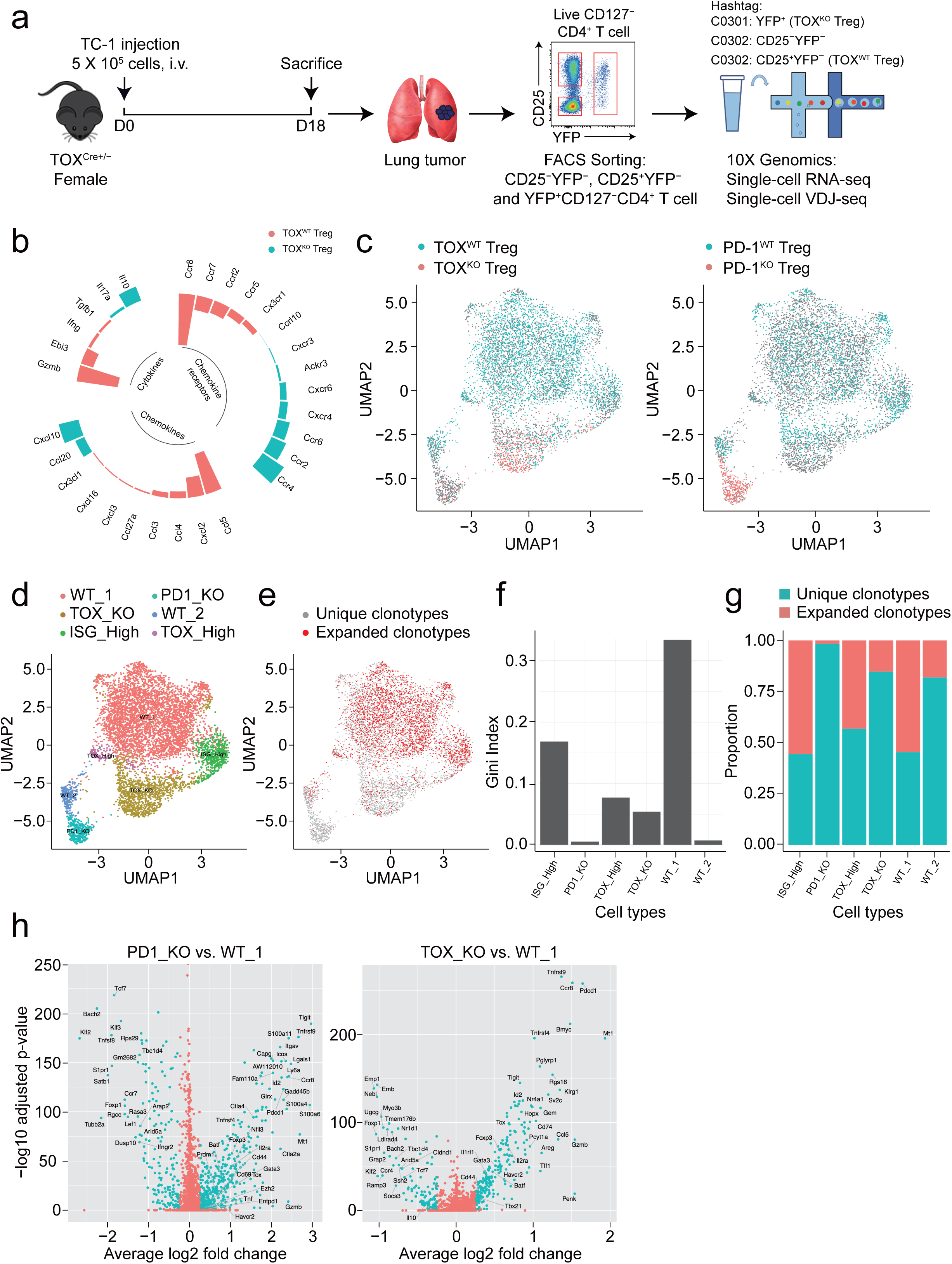
Integrated single-cell RNA and V(D)J sequencing analysis of TOX^WT^, TOX^KO^, PD-1^WT^, and PD-1^KO^ Tregs in the TME (related to Fig. 6). **a**, Experimental workflow for single-cell RNA and V(D)J sequencing. Lung tumors from TC-1 tumor-bearing TOX^Cre+/−^ mice were harvested on day 18 post-injection. Live CD127^−^CD4^+^ T cells were sorted into three populations based on YFP and CD25 expression: YFP^−^CD25^−^, YFP^−^CD25^+^ (TOX^WT^ Tregs), and YFP^+^ (TOX^KO^ Tregs). Sorted populations were labeled with cell hashing antibodies (TotalSeq™-C0301 or C0302) and processed for scRNA- and scV(D)J sequencing using the 10X Genomics Chromium platform. **b**, Circular plot illustrating differentially expressed cytokines, chemokines, and chemokine receptors between TOX^WT^ and TOX^KO^ TI Tregs. Bar height indicates relative expression levels. **c–h**, Integrated analysis of single-cell transcriptomes from PD-1^WT^ and PD-1^KO^ TI Tregs (GSE164033) combined with TOX^WT^ and TOX^KO^ TI Treg datasets. **c**, UMAP visualization of integrated Treg populations, colored by TOX status (left) or PD-1 status (right). **d**, UMAP plot showing six distinct transcriptional clusters: WT_1, WT_2, TOX_KO, PD1_KO, ISG_High, and TOX_High. **e**, UMAP projection displaying the distribution of unique (grey) and expanded (red) T cell receptor clonotypes across integrated Treg populations. **f**, Gini index quantifying the degree of clonal expansion in each Treg subset. Higher values indicate greater clonal dominance. **g**, Stacked bar plot showing the proportion of unique versus expanded clonotypes in each Treg population. **h**, Volcano plots displaying differentially expressed genes between PD-1_KO versus WT Tregs (left) and TOX_KO versus WT Tregs (right). i.v., intravenous; WT, wild type; KO, knock-out; PD-1, programmed death-1; ISG, Interferon-Stimulated Gene.

**Extended Data Fig. 8.**
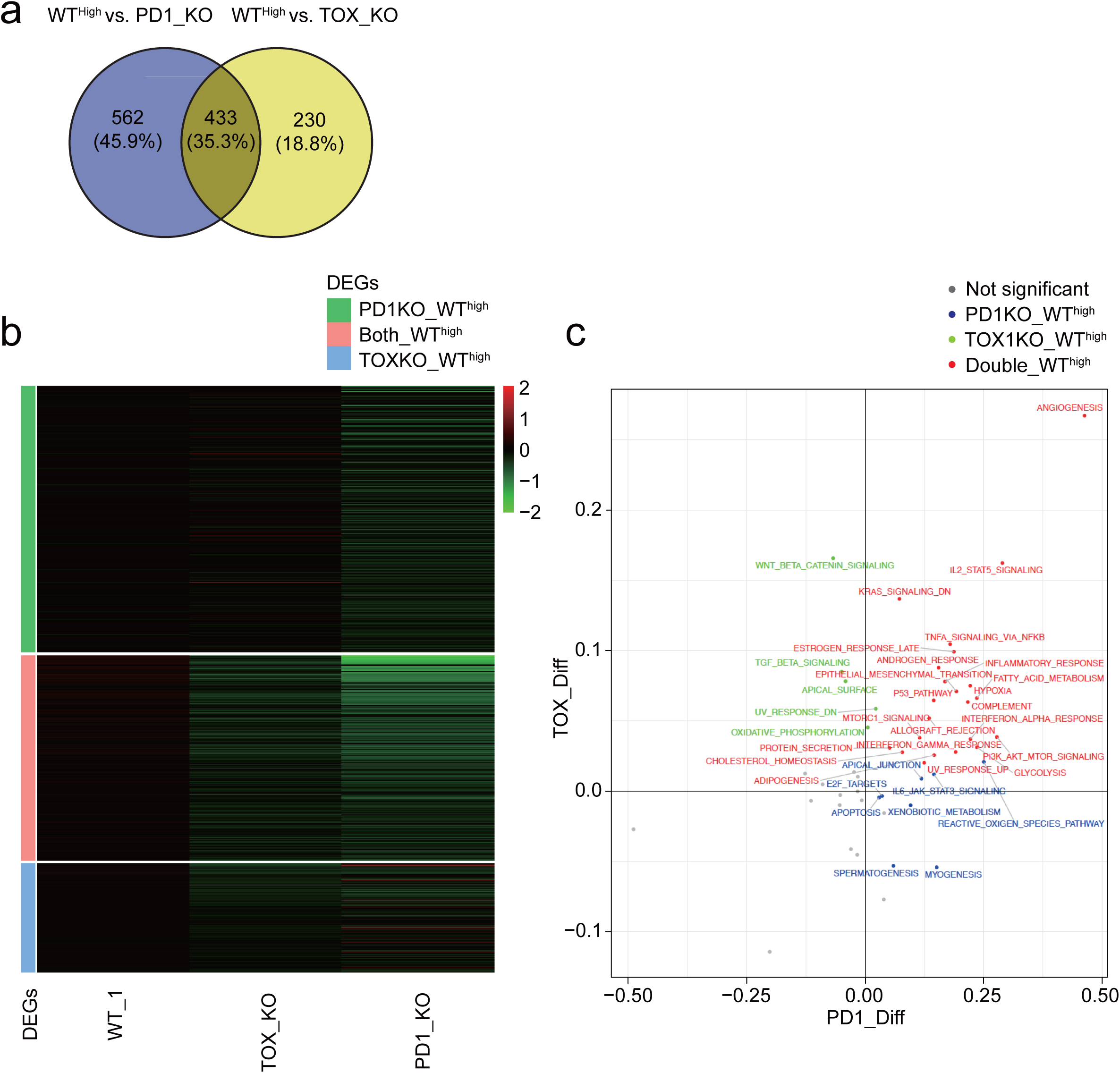
Comparative transcriptomic signatures and pathway enrichment of TOX- and PD-1-deficient Tregs in the TME (related to Fig. 6). **a**, Venn diagram showing the overlap of differentially expressed genes (DEGs) between WT^High^ Tregs versus PD-1^KO^ Tregs and WT^High^ Tregs versus TOX^KO^ Tregs. Numbers indicate unique and shared DEGs, with percentages shown in parentheses. **b**, Heatmap showing the expression patterns of DEGs across three Treg populations: WT^high^, TOX_KO, and PD1_KO as shown in (**a**). Genes are grouped based on their differential expression pattern: genes in PD1_KO versus WT^high^ (green), shared genes in WT^high^ (red), and unique genes in TOX_KO versus WT^high^ (blue). Color scale represents normalized gene expression (z-score), with green indicating low expression and red indicating high expression. DEGs, differentially expressed genes. **c**, Pathway enrichment analysis comparing transcriptional programs altered in each group, visualized across the two axes. Each point represents a biological pathway, with axis values indicating the differential enrichment score (PD1_Diff on x-axis, TOX_Diff on y-axis). Pathways are color-coded: red indicates pathways enriched in WT^high^ Tregs relative to both knockout populations (upper right quadrant), green indicates pathways specifically enriched in TOX KO Tregs (upper left quadrant), blue indicates pathways specifically enriched in PD1 KO Tregs, and grey indicates non-significant pathways.

**Extended Data Fig. 9.**
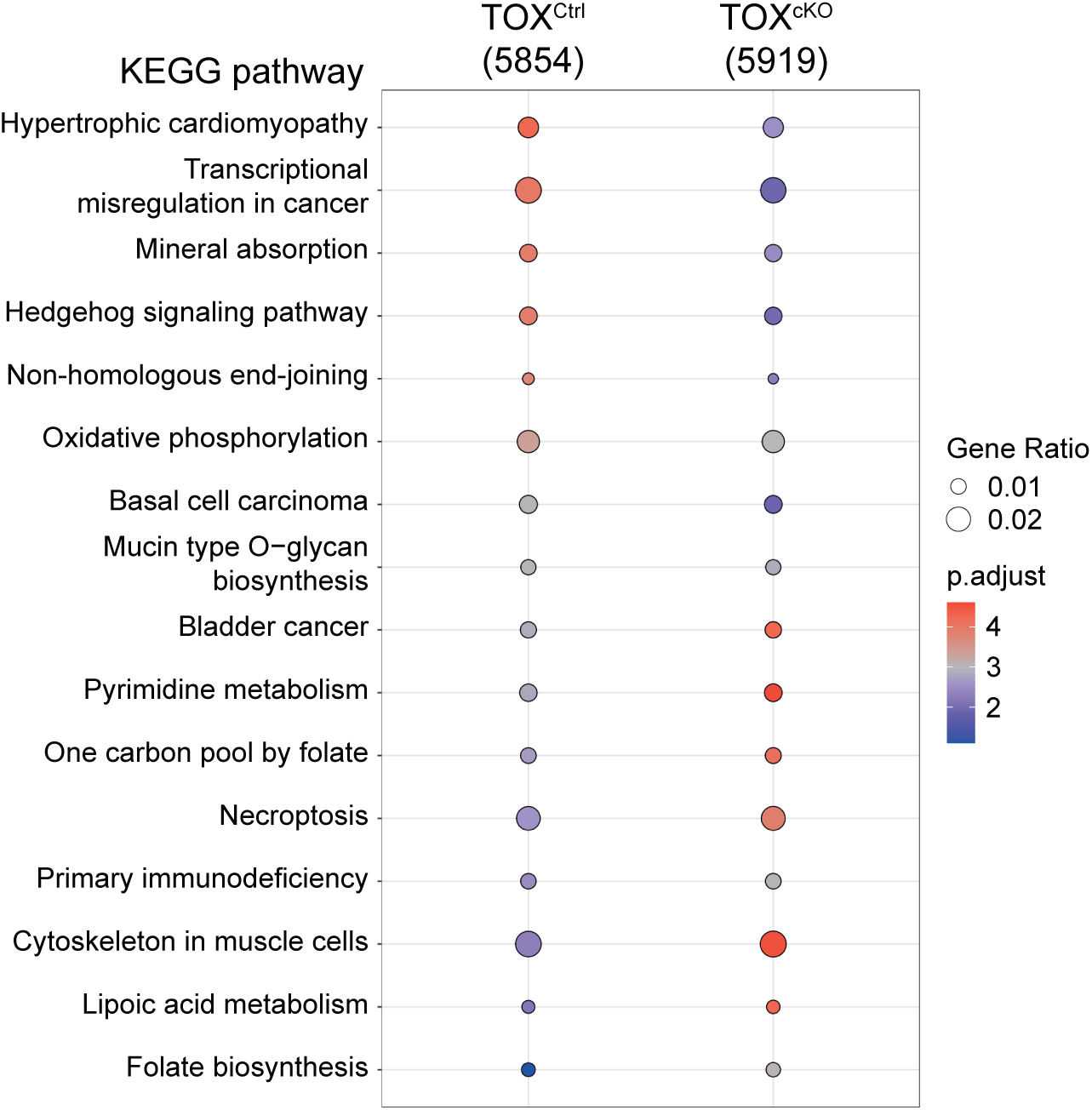
KEGG pathway enrichment analysis of chromatin accessibility in TOX^Ctrl^ and TOX^cKO^ TI Tregs (related to Fig. 7). KEGG pathway analysis comparing open chromatin–associated genes between TOX^Ctrl^ (*n* = 5854 peaks) and TOX^cKO^ (*n* = 5919 peaks) Tregs isolated from TC-1 tumor-bearing mice. Each dot represents a significantly enriched pathway. Dot size indicates the percentage of genes in each pathway, and dot color represents the enrichment score (red indicates higher enrichment, and blue indicates lower enrichment).

**Extended Data Fig. 10.**
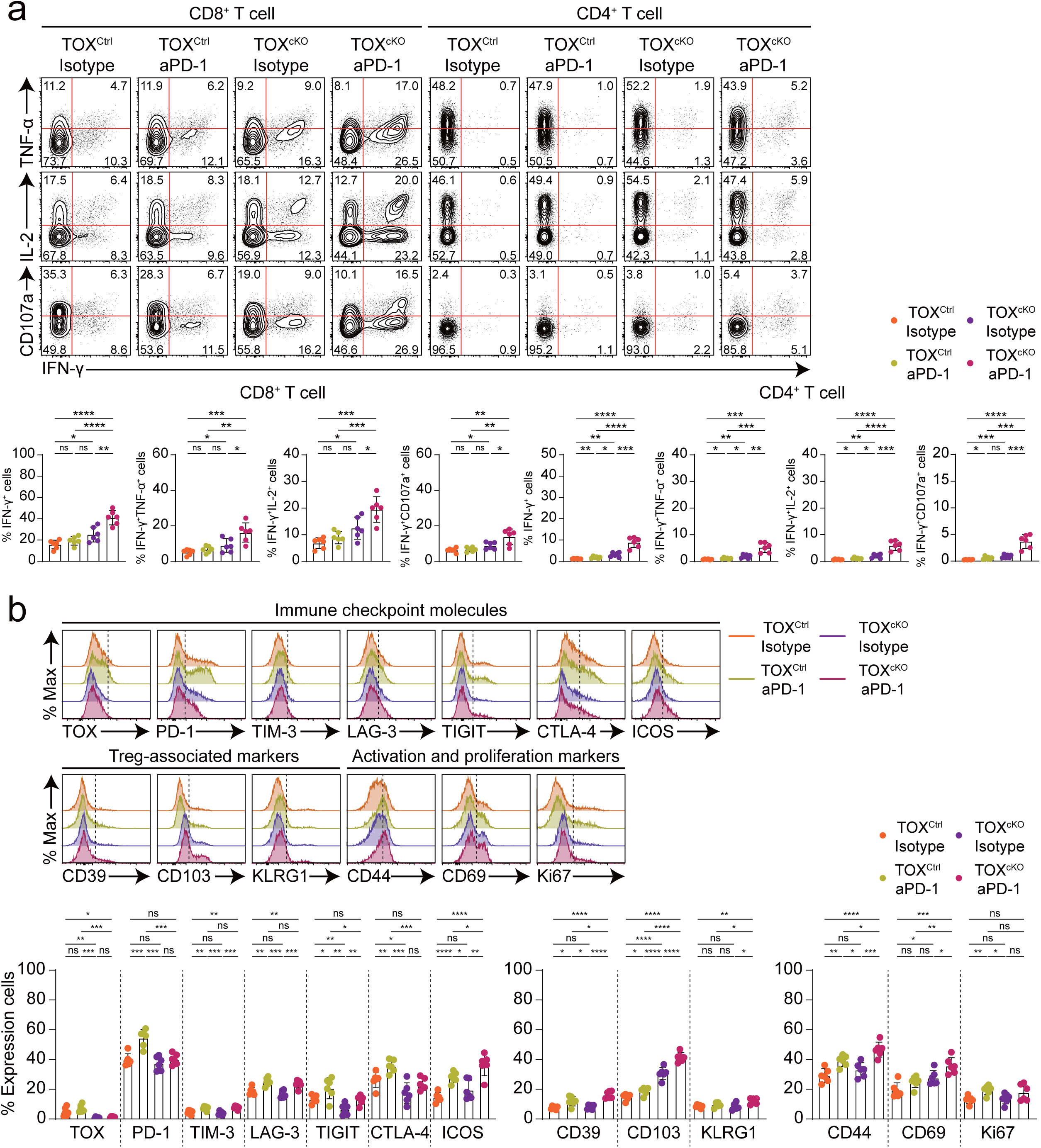
Systemic T cell responses and splenic Treg phenotypes following Treg-specific TOX deficiency and PD-1 blockade (related to Fig. 8). a, Flow cytometric analysis of splenic CD8^+^ and CD4^+^ T cell effector function from each experimental group. Cytokine production was assessed by intracellular staining for IFN-γ, TNF-α, IL-2, and degranulation was assessed by CD107a expression. Representative contour plots (top) and quantitative bar graphs (bottom) show frequencies of IFN-γ^+^, IFN-γ^+^TNF-α^+^, IFN-γ^+^IL-2^+^, and IFN-γ^+^CD107a^+^ cells among CD8^+^ and CD4^+^ T cells. **b**, Flow cytometric analysis of splenic Tregs from the indicated experimental groups. Expression of TOX, immune checkpoint molecules (PD-1, TIM-3, LAG-3, TIGIT, CTLA-4, and ICOS), Treg-associated markers (CD39, CD103, and KLRG1), and activation/proliferation markers (CD44, CD69, and Ki67) was assessed. Histograms (top) depict expression profiles across groups, and bar graphs (bottom) quantify the percentage of positive cells for each marker. Data were concatenated within each group (**a, b**). The numbers in the plots indicate the percentage of the population (**a**). All bar graphs show mean ± SEM. Statistical analysis was performed using two-tailed unpaired Student’s *t*-test. ns, not significant; *, *P* < 0.05; **, *P* < 0.01; ***, *P* < 0.001; ****, *P* < 0.0001.

## REFERENCES

1. Josefowicz, S.Z., Lu, L.F. & Rudensky, A.Y. Regulatory T cells: mechanisms of differentiation and function. Annu Rev Immunol 30, 531–564 (2012).

2. Lee, W. & Lee, G.R. Transcriptional regulation and development of regulatory T cells. Exp Mol Med 50, e456 (2018).

3. Scott, E.N., Gocher, A.M., Workman, C.J. & Vignali, D.A.A. Regulatory T Cells: Barriers of Immune Infiltration Into the Tumor Microenvironment. Front Immunol 12, 702726 (2021).

4. Li, C., Jiang, P., Wei, S., Xu, X. & Wang, J. Regulatory T cells in tumor microenvironment: new mechanisms, potential therapeutic strategies and future prospects. Mol Cancer 19, 116 (2020).

5. Yang, J. & Bae, H. Drug conjugates for targeting regulatory T cells in the tumor microenvironment: guided missiles for cancer treatment. Exp Mol Med 55, 1996–2004 (2023).

6. Kim, J.H., Kim, B.S. & Lee, S.K. Regulatory T Cells in Tumor Microenvironment and Approach for Anticancer Immunotherapy. Immune Netw 20, e4 (2020).

7. Tay, C., Tanaka, A. & Sakaguchi, S. Tumor-infiltrating regulatory T cells as targets of cancer immunotherapy. Cancer Cell 41, 450–465 (2023).

8. Kang, J.H. & Zappasodi, R. Modulating Treg stability to improve cancer immunotherapy. Trends Cancer 9, 911–927 (2023).

9. Plitas, G. et al. Regulatory T Cells Exhibit Distinct Features in Human Breast Cancer. Immunity 45, 1122–1134 (2016).

10. De Simone, M. et al. Transcriptional Landscape of Human Tissue Lymphocytes Unveils Uniqueness of Tumor-Infiltrating T Regulatory Cells. Immunity 45, 1135–1147 (2016).

11. Zheng, L. et al. Pan-cancer single-cell landscape of tumor-infiltrating T cells. Science 374, abe6474 (2021).

12. Kim, M.J. et al. Deletion of PD-1 destabilizes the lineage identity and metabolic fitness of tumor-infiltrating regulatory T cells. Nat Immunol 24, 148–161 (2023).

13. Magnuson, A.M. et al. Identification and validation of a tumor-infiltrating Treg transcriptional signature conserved across species and tumor types. Proc Natl Acad Sci U S A 115, E10672–E10681 (2018).

14. Wertheimer, T. et al. IL-23 stabilizes an effector T(reg) cell program in the tumor microenvironment. Nat Immunol 25, 512–524 (2024).

15. Itahashi, K., et al. BATF epigenetically and transcriptionally controls the activation program of regulatory T cells in human tumors. Sci Immunol 7, eabk0957 (2022).

16. Shan, F., et al. Integrated BATF transcriptional network regulates suppressive intratumoral regulatory T cells. Sci Immunol 8, eadf6717 (2023).

17. Cretney, E. et al. The transcription factors Blimp-1 and IRF4 jointly control the differentiation and function of effector regulatory T cells. Nat Immunol 12, 304–311 (2011).

18. Shi, L.Z. et al. HIF1alpha-dependent glycolytic pathway orchestrates a metabolic checkpoint for the differentiation of TH17 and Treg cells. J Exp Med 208, 1367–1376 (2011).

19. Dang, E.V. et al. Control of T(H)17/T(reg) balance by hypoxia-inducible factor 1. Cell 146, 772–784 (2011).

20. Niu, H. & Wang, H. TOX regulates T lymphocytes differentiation and its function in tumor. Front Immunol 14, 990419 (2023).

21. O’Flaherty, E. & Kaye, J. TOX defines a conserved subfamily of HMG-box proteins. BMC Genomics 4, 13 (2003).

22. Wilkinson, B. et al. TOX: an HMG box protein implicated in the regulation of thymocyte selection. Nat Immunol 3, 272–280 (2002).

23. Aliahmad, P. & Kaye, J. Development of all CD4 T lineages requires nuclear factor TOX. J Exp Med 205, 245–256 (2008).

24. Aliahmad, P., Kadavallore, A., de la Torre, B., Kappes, D. & Kaye, J. TOX is required for development of the CD4 T cell lineage gene program. J Immunol 187, 5931–5940 (2011).

25. Seehus, C.R. et al. The development of innate lymphoid cells requires TOX-dependent generation of a common innate lymphoid cell progenitor. Nat Immunol 16, 599–608 (2015).

26. Xu, W. et al. The Transcription Factor Tox2 Drives T Follicular Helper Cell Development via Regulating Chromatin Accessibility. Immunity 51, 826–839 e825 (2019).

27. Aliahmad, P., de la Torre, B. & Kaye, J. Shared dependence on the DNA-binding factor TOX for the development of lymphoid tissue-inducer cell and NK cell lineages. Nat Immunol 11, 945–952 (2010).

28. Beltra, J.C. et al. Developmental Relationships of Four Exhausted CD8(+) T Cell Subsets Reveals Underlying Transcriptional and Epigenetic Landscape Control Mechanisms. Immunity 52, 825–841 e828 (2020).

29. Khan, O. et al. TOX transcriptionally and epigenetically programs CD8(+) T cell exhaustion. Nature 571, 211–218 (2019).

30. Kim, K. et al. Single-cell transcriptome analysis reveals TOX as a promoting factor for T cell exhaustion and a predictor for anti-PD-1 responses in human cancer. Genome Med 12, 22 (2020).

31. Scott, A.C. et al. TOX is a critical regulator of tumour-specific T cell differentiation. Nature 571, 270–274 (2019).

32. Wang, X. et al. TOX promotes the exhaustion of antitumor CD8(+) T cells by preventing PD1 degradation in hepatocellular carcinoma. J Hepatol 71, 731–741 (2019).

33. Chen, J. et al. Single-cell transcriptomics reveal the intratumoral landscape of infiltrated T-cell subpopulations in oral squamous cell carcinoma. Mol Oncol 15, 866–886 (2021).

34. Zhao, Y. et al. Increased TOX expression associates with exhausted T cells in patients with multiple myeloma. Exp Hematol Oncol 11, 12 (2022).

35. Seo, H. et al. TOX and TOX2 transcription factors cooperate with NR4A transcription factors to impose CD8(+) T cell exhaustion. Proc Natl Acad Sci U S A 116, 12410–12415 (2019).

36. Kim, C.G., et al. VEGF-A drives TOX-dependent T cell exhaustion in anti-PD-1-resistant microsatellite stable colorectal cancers. Sci Immunol 4 (2019).

37. Huang, S. et al. Increased TOX expression concurrent with PD-1, Tim-3, and CD244 in T cells from patients with non-Hodgkin lymphoma. Asia Pac J Clin Oncol 18, 143–149 (2022).

38. Lamarche, C. et al. Tonic-signaling chimeric antigen receptors drive human regulatory T cell exhaustion. Proc Natl Acad Sci U S A 120, e2219086120 (2023).

39. Yu, G., Wang, L.G., Han, Y. & He, Q.Y. clusterProfiler: an R package for comparing biological themes among gene clusters. OMICS 16, 284–287 (2012).

40. Liberzon, A. et al. The Molecular Signatures Database (MSigDB) hallmark gene set collection. Cell Syst 1, 417–425 (2015).

41. Liberzon, A. et al. Molecular signatures database (MSigDB) 3.0. Bioinformatics 27, 1739–1740 (2011).

42. Wang, H. et al. CD36-mediated metabolic adaptation supports regulatory T cell survival and function in tumors. Nat Immunol 21, 298–308 (2020).

43. Stuart, T. et al. Comprehensive Integration of Single-Cell Data. Cell 177, 1888–1902 e1821 (2019).

44. Hanzelmann, S., Castelo, R. & Guinney, J. GSVA: gene set variation analysis for microarray and RNA-seq data. BMC Bioinformatics 14, 7 (2013).

45. Zhang, Y. et al. Model-based analysis of ChIP-Seq (MACS). Genome Biol 9, R137 (2008).

46. Amemiya, H.M., Kundaje, A. & Boyle, A.P. The ENCODE Blacklist: Identification of Problematic Regions of the Genome. Sci Rep 9, 9354 (2019).

47. Yu, G., Wang, L.G. & He, Q.Y. ChIPseeker: an R/Bioconductor package for ChIP peak annotation, comparison and visualization. Bioinformatics 31, 2382–2383 (2015).

48. Lawrence, M. et al. Software for computing and annotating genomic ranges. PLoS Comput Biol 9, e1003118 (2013).

49. Kanehisa, M., Furumichi, M., Sato, Y., Kawashima, M. & Ishiguro-Watanabe, M. KEGG for taxonomy-based analysis of pathways and genomes. Nucleic Acids Res 51, D587–D592 (2023).

50. Ou, J. & Zhu, L.J. trackViewer: a Bioconductor package for interactive and integrative visualization of multi-omics data. Nat Methods 16, 453–454 (2019).

51. Bentsen, M. et al. ATAC-seq footprinting unravels kinetics of transcription factor binding during zygotic genome activation. Nat Commun 11, 4267 (2020).

52. Gao, Y. et al. Intratumoral stem-like CCR4+ regulatory T cells orchestrate the immunosuppressive microenvironment in HCC associated with hepatitis B. J Hepatol 76, 148–159 (2022).

53. Naizir, B. et al. TOX drives CD4(+) T(H)1 effector function, antitumor immunity and autoimmune pathology. Nat Immunol 27, 1013–1025 (2026).

