## Supplementary materials for "TOX enforces the immunosuppressive program of tumor-infiltrating regulatory T cells"

### Contents

**Supplementary Table 1.** Baseline characteristics of HNSCC patients

**Supplementary Table 2.** Baseline characteristics of NSCLC patients

**Supplementary Figure.** Graphical abstract

**Supplementary Table 1.** Baseline characteristics of HNSCC patients

| **Characteristics** | **Number (%)** |
|---|---|
| Total number of patients | 23 |
| *Age* in years (median, range) | 58 (38-72) |
| *Sex* | |
| Male | 21 (91.3%) |
| Female | 2 (8.7%) |
| *Cancer type* | |
| Laryngeal cancer | 1 (4.4%) |
| Oral cavity cancer | 3 (13.0%) |
| Oropharyngeal cancer | 19 (82.6%) |
| *Human papilloma virus infection* | |
| Positive | 18 (78.3%) |
| Negative | 5 (21.7%) |
| *Stage* | |
| I | 5 (21.8%) |
| II | 7 (30.4%) |
| III | 7 (30.4%) |
| IV | 4 (17.4%) |

**Supplementary Table 2.** Baseline characteristics of NSCLC patients

| Characteristics | Number (%) |
|---|---|
| Total number of patients | 35 |
| *Age* in years (median, range) | 67 (22-83) |
| *Sex* | |
| Male | 20 (57.1%) |
| Female | 15 (42.9%) |
| *Smoking History* | |
| Never | 18 (51.4%) |
| Former | 14 (40%) |
| Current | 3 (8.6%) |
| *Cancer type* | |
| Adenocarcinoma | 34 (97.1%) |
| Squamous cell carcinoma | 1 (2.9%) |
| *PD-L1* | |
| Positive (≥1%) | 12 (34.3%) |
| Negative | 23 (65.7%) |
| *Stage* | |
| I | 21 (60%) |
| II | 6 (17.1%) |
| III | 8 (22.9%) |

**Supplementary Figure.** Graphical abstract

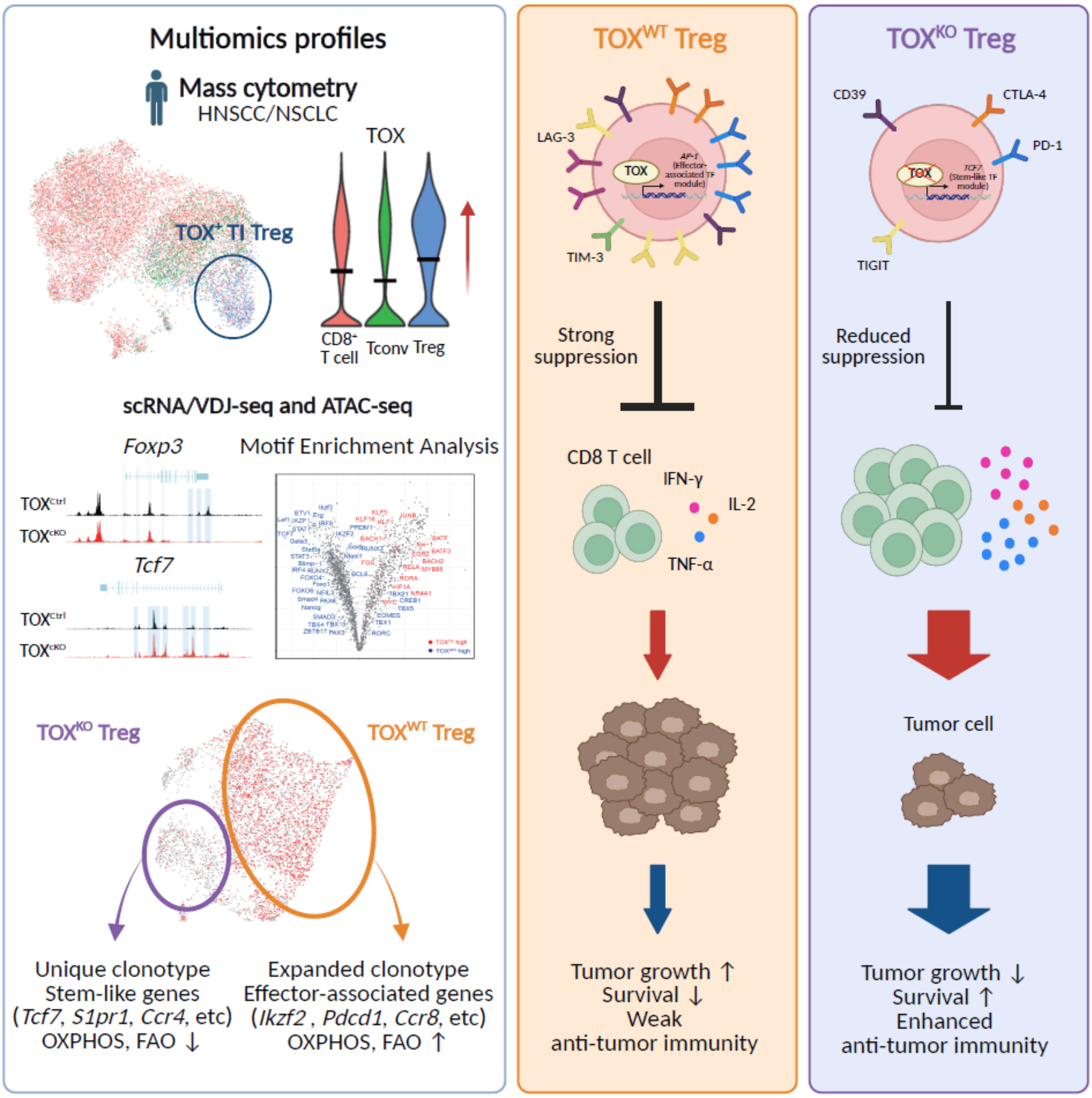
